# Investigating the Evolution of Green Algae with a Large Transcriptomic Dataset

**DOI:** 10.1101/2024.02.21.581324

**Authors:** David A. Ferranti, Charles F. Delwiche

**Affiliations:** Cell Biology and Molecular Genetics, Biosciences Research Building, University of Maryland College Park, College Park, MD 20742, USA; Ecology, Behavior, and Evolution, Muir Biology Building, University of California San Diego, La Jolla, CA 92093, US

## Abstract

The colonization of land by plants, thought to have occurred approximately 450-500 million years ago (Ma) is one of the most important events in the history of life on Earth. Land plants, hereafter referred to as “embryophytes,” comprise the foundation of every terrestrial biome, making them an essential lineage for the origin and maintenance of biodiversity. The embryophytes form a monophyletic clade within one of the two major phyla of the green algae, the Streptophyta. Estimates from fossil data and molecular clock analyses suggest the charophytes diverged from the other main phylum of green algae, the Chlorophyta, as much as 1500 Ma. Here we present a phylogenetic analysis using transcriptomic and genomic data of 62 green algae and embryophyte operational taxonomic units, 31 of which were assembled *de novo* for this project. We focus on identifying the charophyte lineage that is sister to embryophytes, and show that the Zygnematophyceae have the strongest support, followed by the Charophyceae. We demonstrate that this phylogenetic tree topology is robust across different phylogenetic models and methods. Furthermore, we examine amino acid and codon usage across the tree and find these data broadly follow the phylogenetic tree. We conclude by searching the dataset for several protein domains and gene families known to be important in embryophytes, including the ethylene signaling pathway and various ion transporters. Many of these domains and genes have homologous sequences in the charophyte lineages, giving insight into the processes that underlay the colonization of the land by plants.

## Introduction

The colonization of terrestrial habitats by land plants roughly 500 Ma is one of the most important events in the history of complex life. Land plants, hereafter referred to as “embryophytes” comprise the ecological foundation of every major biome (the term “embryophyte” refers to their characteristic life cycle involving the alternation of multicellular haploid and diploid generations; Heckman *et al*. 2001; Campbell 1940). When embryophytes emerged in the fossil record there was a profound restructuring of the entire biosphere, with a substantial increase in burial of carbon leading to an increase in atmospheric oxygen and deposition of coal, alteration of erosional and hydrological processes as well as formation of mud and mudstone, and substantial alteration in marine chemistry stemming from the changed composition of runoff (D’Antonio *et al*. 2019; McMahon *et al*. 2018; Wellman and Strother 2015; Lyons *et al*. 2014). Embryophytes are also foundational crop species and thus critical to the development of complex human societies. Additionally, many embryophytes produce other important products for global commerce or have medicinal applications. The embryophytes form a monophyletic clade within one of the two major phyla of the green algae, the Streptophyta, or “Streptophytes” (sensu lato) (Delwiche and Cooper 2015; Delwiche and Timme 2011; Mattox and Stewart 1984). The other major phylum, the Chlorophyta, contains the bulk of green algal diversity and nearly all of its marine representatives, while a third small lineage (Palmophyllophyta, or Prasinodermophyta) comprising *Verdigellas*, *Palmophyllum*, *Prasinococcus* and *Prasinoderma* may be an outgroup to both Streptophya and Chlorophyta (Li *et al*. 2020). For the purposes of this manuscript, Streptophytes refers to the entire charophyte lineage and all its descendants, including embryophytes, whereas “charophytes” refers to only the charophyte algae, excluding embryophytes. “Charophytes s. s.” (sensu stricto) refers only to the lineage including *Chara, Nitella, Tolypella*, and the other organisms known informally as stoneworts; we also add “s.l” as needed for clarity. It should be noted that the fossil diversity of charophytes s.s. greatly exceeds their extant diversity, but that only a minimal fossil record exists for other non-embryophyte charophytes (Tappan 1980).

Estimates from both fossil data and molecular clock analyses suggest that the Streptophytes diverged from the other main phylum of green algae, the “chlorophytes,” by as much as 1500 Ma (Del Cortona *et al*. 2020; Tang *et al*. 2020; Leliaert *et al*. 2011; Hedges *et al*. 2004). Together, the Streptophytes and the chlorophytes (and perhaps a few additional taxa such as *Prasinoderma*) comprise the “Chloroplastida,” or green plants (Adl *et al*. 2019). Charophyte algae consist entirely of freshwater organisms, although a few taxa have adapted to persist in subaerial and saline environments, whereas chlorophytes, although likely freshwater in origin, are found in marine, freshwater, and terrestrial environments, and independently colonized land several times (Fučíková *et al*. 2014; Blank 2013; Leliaert *et al*. 2012). Most studies focused on characterizing chlorophyte evolutionary relationships have dealt with the “core chlorophyte” clades (Chlorophyceae, Trebouxiophyceae, and Ulvophyceae), although there have been efforts in recent years to examine the more basal chlorophytes, including the deep-water marine Palmophyllophyceae and the various “prasinophyte” (scaly flagellate) clades (Li *et al*. 2021; Li *et al*. 2020; Leliaert *et al*. 2016). The Trebouxiophyceae are typically thought to be the outgroup to the remaining two core chlorophyte clades, although it has been proposed that neither Trebouxiophyceae nor Ulvophyceae are truly monophyletic, and a new classification system is needed to properly describe the core chlorophyte clades (Del Cortona *et al*. 2020; Fučíková *et al*. 2014). More than twenty thousand species of green algae have been described so far, although the true number is likely higher (Christenhusz *et al*. 2016; Guiry 2012). Much of the work on the evolution of the Streptophytes relates to the placement of different charophyte lineages in relationship to each other and to modern land plants. In particular, the identification of the specific charophyte s. l. lineage that is the sister to embryophytes has been studied for decades (Lewis and McCourt 2004; Turmel *et al*. 2003; Karol *et al*. 2001; Bhattacharya and Medlin 1998; Graham 1996; Mishler and Churchill 1985). Understanding the evolutionary relationship between the Streptophytes and embryophytes and the features that allowed for the algal transition onto land is important to advancing our knowledge of how embryophytes came to dominate terrestrial ecosystems (Bowles *et al*. 2020; Cheng *et al*. 2019; Harholt *et al*. 2016).

Three separate charophyte algal clades are prominent candidates for the sister taxon to embryophytes: the Charophyceae s. s., the Coleochaetophyceae, and the Zygnematophyceae. Taxa in all three clades display complex morphology, multicellularity, and sexual reproduction, with zygnematophytes undergoing conjugation. Prior to the advent of molecular phylogenetics, these features established the Charophyceae s. s., the Coleochaetophyceae, and the Zygnematophyceae as the primary candidates under consideration for the immediate outgroup to embryophytes (Wicket *et al*. 2014; Graham 1993). Since then, separate phylogenetic analyses using a combination of plastid, mitochondrial, and nuclear genes have recovered support for each clade being the true sister taxon (Glass *et al*. 2023; Finet *et al*. 2010; but see Laurin-Lemay *et al*. 2012; Turmel *et al*. 2006; McCourt *et al*. 2004; Turmel *et al*. 2003; Karol *et al*. 2001; Mishler *et al*. 1994). Much of the diversity of Zygnematophyceae is within a single family, the Desmidiaceae, which is a derived clade consisting of single-celled and chain-forming species. Single-celled zygnematophytes are informally known as desmids whether or not they are in the Desmidiaceae, and there is a complex distribution of uni- and multi- cellularity within the Zygnematophyceae (Cheng *et al*. 2019; Gontcharov and Melkonian 2011; Hall *et al*. 2008). The distribution of multicellularity within the Streptophyta implies that the ancestor of the three “higher charophyte” clades was multicellular and thereafter lost one or more times in the zygnematophytes (Delwiche and Cooper 2015; but see Cheng *et al*. 2019; Wicket *et al*. 2014). Although the most current phylogenomic studies indicate that the Zygnematophyceae are the sister lineage to embryophytes, additional work with denser taxon sampling from the different green algal lineages is needed to resolve the topology of the Chloroplastida phylogenetic tree (Wickett *et al*. 2014; Timme *et al*. 2012). Previous analyses have significantly advanced our understanding of the transition of the charophyte algae onto land, but often relied on either only a few algal taxa or a handful of genes.

The advent of high-throughput sequencing data has allowed for the development of many high-quality omics resources for many non-model organisms, green algae and early-diverging embryophytes among them (Jiao *et al*. 2020; Cheng *et al*. 2019; Nishiyama *et al*. 2018; Delwiche *et al*. 2017; Hori *et al*. 2014). Both transcriptomic and genomic data are powerful in constructing phylogenies using multiple sequence alignments from dozens or even hundreds of genes. These large alignments are key to reconstructing deep evolutionary relationships, as they give sufficient power for complex phylogenetic construction and the testing of evolutionary hypotheses (Cheon *et al*. 2020; McKain *et al*. 2018; Wickett *et al*. 2014). Due to the high biological diversity and long divergence times found across Chloroplastida, it has been difficult to answer questions that span the evolution of the entire clade. Even identifying orthologous sequences from across the tree for alignment and additional analysis can be difficult because some clades, especially embryophytes, have undergone rapid genome evolution (Proost *et al*. 2011). Furthermore, phylogenetic analysis may become inconsistent and generate artifacts when highly diverged taxa are included, and polygenic datasets only exacerbate this problem (Hillis *et al*. 1992; Felsenstein 1978). Long branch attraction is therefore more likely to occur when analyzing phylogenies that span hundreds of millions of years and have been strongly pruned by extinction (Brinkmann *et al*. 2005). Recent advances, including application of models for heterotachous sequence evolution, and increased breadth of taxon sampling, have provided valuable new insights into the diversification of Chloroplastida, but many unsolved problems remain (Glass *et al*. 2023; Del Cortona *et al*. 2020; Cox *et al*. 2014; Zhong *et al*. 2014; Rodríguez-Ezpeleta *et al*. 2007).

The sequencing and assembly of several *Chara* genomes, as well as the genomes of several species of Zygnematophyceae, have provided valuable resources for investigation of the sister lineage to embryophytes with different phylogenetic methods. Furthermore, both bioinformatic analyses and manipulative laboratory experiments performed on various Zygnematophyceae taxa have shown groups of orthologous genes associated with stress in embryophytes as well as similar cellular responses to desiccation and intense light (Jiao *et al*. 2020; Li *et al*. 2020; Cheng *et al*. 2019; De Vries *et al*. 2018; Holzinger and Pichrtová 2017). It has been suggested that the charophyte algae were uniquely suited to survive in terrestrial habitats due to their genomic and molecular toolkit, although it seems that a complex combination of attributes was required (De Vries *et al*. 2018; Delaux *et al*. 2015). Mapping the presence of different components of a modern-day organism’s genomic toolkit throughout a phylogenetic tree is a useful method for generating hypotheses about the emergence of important biological pathways.

Here we present a maximum-likelihood phylogenetic analysis using an alignment built from more than one thousand three hundred orthologous sequences of 62 operational taxonomic units that span the Chloroplastida tree of life from early-diverging marine prasinophytes (single-celled flagellate green algae, but not a monophyletic group) to angiosperms. We focus on identifying the charophyte algal lineage that is sister to embryophytes by testing a series of alternative phylogenetic tree topologies that shuffle the positions of the Zygnematophyceae, Coleochaetophyceae, and Charophyceae s. s. clades in relation to both embryophytes and the older green algae clades. We further contextualize the phylogenetic analyses by examining how both amino acid and codon usage vary across the phylogenetic tree. We conclude by tracing the presence of several protein domains found in the ethylene signaling pathway as well as several ion transporter gene families through the different algal lineages.

## Methods

### *De novo* assemblies and dataset construction

We generated 32 paired-end RNA-Seq datasets by extracting poly-A selected mRNA from 31 species of green algae and sequenced it high-throughput shotgun sequencing on an Illumina machine (Table S1). Two libraries were prepared for *Nitella mirabilis,* one each from the upper and lower portions of the organism. A single library was prepared for all other taxa. Reads were de-multiplexed using the Illumina Casava pipeline. To remove poor-quality reads, poly-A tails, and singletons, reads were then lightly filtered with PRINSEQ-lite version 0.20.4 running on the following settings: -trim_qual_left 20, -trim_qual_right 20, -trim_qual_window 20, - trim_tail_left 101, -trim_tail_right 101, -trim_ns_left 1, -trim_ns_right 1, -min_len 25, - min_qual_mean 20, and -out_format 3 (Schmieder and Edwards 2011).

This set of filtered reads was used for all subsequent analyses. The filtered reads were assembled using rnaSPAdes 3.14.0 with default options (Bushmanova *et al*. 2019). In the case of *Nitella mirabilis*, both pairs of reads, from the libraries prepared from the upper and lower parts of the plant, were combined to assemble the transcriptome. Assemblies were converted into amino acid sequences with TransDecoder v5.5.0 and assessed for completeness by running BUSCO v4.0.6 in protein mode against the Chlorophyte, Viridiplantae, and Eukaryote databases (Seppey *et al*. 2019; Haas *et al*. 2013).

Data for other taxa were either downloaded from publicly available sources or provided by collaborators as transcriptomes or coding sequences from the genome (Table S2). For both types of data, the nucleotide sequences were converted into amino acid sequences with TransDecoder v5.5.0 and assessed for completeness using BUSCO as with the *de novo* transcriptome assemblies described above.

### Ortholog identification and multiple sequence alignment

Because traditional reciprocal BLAST search methods found relatively few orthologous genes, we used a hidden Markov model approach to identify orthologs. First, coding sequences were obtained from nine (three charophyte and six chlorophyte) different green algae genome assemblies (Table S3). The coding sequences were checked for completeness with BUSCO and then converted into amino acid sequences with TransDecoder. Orthofinder was used to identify orthogroups in those nine algae (Emms and Kelly 2018; Emms and Kelly 2015). Orthogroups that contained at least one sequence from each of the nine algae species (hereafter referred to as universal orthogroups) were aligned using MAFFT v7.471 and used to construct hidden Markov models using HMMER3.1 (Eddy 2009; Katoh *et al*. 2002). Orthologs were then identified in each of the species in the larger dataset by searching the species’ assembly with the hidden Markov model generated from each universal orthogroup and extracting the sequence that produced the best hit according to bitscore. The *Chara braunii* and *Penium margaritaceum* genomes were also used in the later phylogenetic analyses.

### Phylogenetic analyses

Individual orthologs were aligned using MAFFT v7.471 and then concatenated into a single superalignment. To reduce the percentage of gaps in the alignment, trimAl v1.3 was used to remove sites in the superalignment that contained a gap for 25% or more taxa (Capella-Gutiérrez *et al*. 2009). IQ-TREE v1.6.12 was used to find a maximum likelihood phylogeny, with the preferred model being the LG substitution matrix, empirical base frequencies, and ten rate parameters (LG+F+R10). Bootstrapping was performed with ten thousand ultrafast bootstrap replicates (Hoang *et al*. 2017; Nguyen *et al*. 2015). The model was selected by using IQ-TREE’s ModelFinder option, which calculates the Bayesian Information Criterion (BIC) for each of 78 total protein models using the JTT, LG, and WAG substitution matrices and up to ten rate parameters (Kalyaanamoorthy *et al*. 2017; Le and Gascuel 2008; Whelan and Goldman 2001; Yang 1995; Jones *et al*. 1992). An additional analysis was performed using the GHOST model with four categories (LG+F*H4) to account for heterotachous sequence evolution (Crotty *et al*. 2020). ModelFinder was used to determine the optimal amino acid matrix (JTT, LG, or WAG) and number of rate parameters for each of the individual trimAl filtered alignments for the 1323 orthologous genes. This was used to perform an edge-unlinked partitioned analysis with each individual gene using its optimal model. The same individual gene trees calculated from the analysis of each individual orthologous gene were fed into ASTRAL III to produce a reconciliation tree (Zhang *et al*. 2018).

We further investigated the topology of the phylogenetic tree by using likelihood mapping analysis and the groupings shown in Table S4 (Strimmer and Von Haeseler 1997). We performed a total of seven likelihood mapping analyses. Three of the analyses ignored the “Older Taxa” grouping and calculated the support under the LG+F+R10 model, the edge-unlinked partition model, and the GHOST heterotachy model. The other four likelihood mapping analyses were performed with the LG+F+R10 model and moved the Older Taxa grouping to be included in each of the Charophyceae s. s., Coleochaetophyceae, Zygnematophyceae, and Embryophyte groupings. In each analysis, we recorded the percentage of likelihood quartets that showed each of the three charophyte algae clades of interest as the sister taxa of Embryophytes in the resulting four-taxon tree. We also calculated the likelihood (under the LG+F+R10 model) of each of the 15 possible 5-taxon tree topologies consisting of the separate Embryophyte, Charophyceae s. s., Coleochaetophyceae, Zygnematophyceae, and Older Taxa groupings.

### Amino acid and codon composition analyses

Amino acid and codon usage were calculated and analyzed via principal components analysis using the FactoMineR and factoextra packages in R (Lê *et al*. 2008; Kassambara 2016).

### Protein domain and protein family searches

We searched for a variety of specific protein domains found in genes involved in the ethylene signaling pathway in land plants, as well as three other gene families. Searches were performed in DIAMOND BLAST with the amino acid sequence of each domain from the corresponding *Arabidopsis thaliana* gene as the query against the combined genomic and transcriptomic dataset included in the ortholog identification and phylogenetic analyses (Buchfink *et al*. 2014). Hits to *Arabidopsis thaliana* were excluded to avoid distortion of the results.

## Results

### Dataset quality and completeness

Taxa are organized by the “Major Clade” classification given in Table S3. Mean BUSCO completeness scores are given in the supplementary data for sequences used for phylogenetic analysis (Figure S1) and HMM construction (Figure S2). OrthoFinder identified a total of 2619 universal orthogroups from the nine algal genomes. After hidden Markov model construction, 1327 of these orthogroups produced a hit for all taxa used in the phylogenetic analysis. The presence/absence of each universal orthogroup in each taxa is shown in Figure 1. After concatenation, TrimAl filtering reduced this number to a final total of 1323 orthologous genes and 566428 sites. The percentage of gap characters for each species included in the phylogenetic analysis after the trimAl filtration step is shown in Figure 2. Both the *Nothoceros* taxa (hornworts) and several gymnosperm taxa show a notably high percentage of gaps compared to the other taxa in the alignment. No green algae species is composed of greater than 20 % gaps, even those with relatively low BUSCO scores. In contrast, among embryophytes, taxa with low BUSCO scores have a high percentage of gaps, perhaps because the HMMs were built from algal sequences. Another source of bias comes from the fact that only genomes from Chlorophyceae were available for HMM construction, with no representatives of the Trebouxiophyceae or Ulvophyceae being available at the time of analysis.

**Figure 1:**
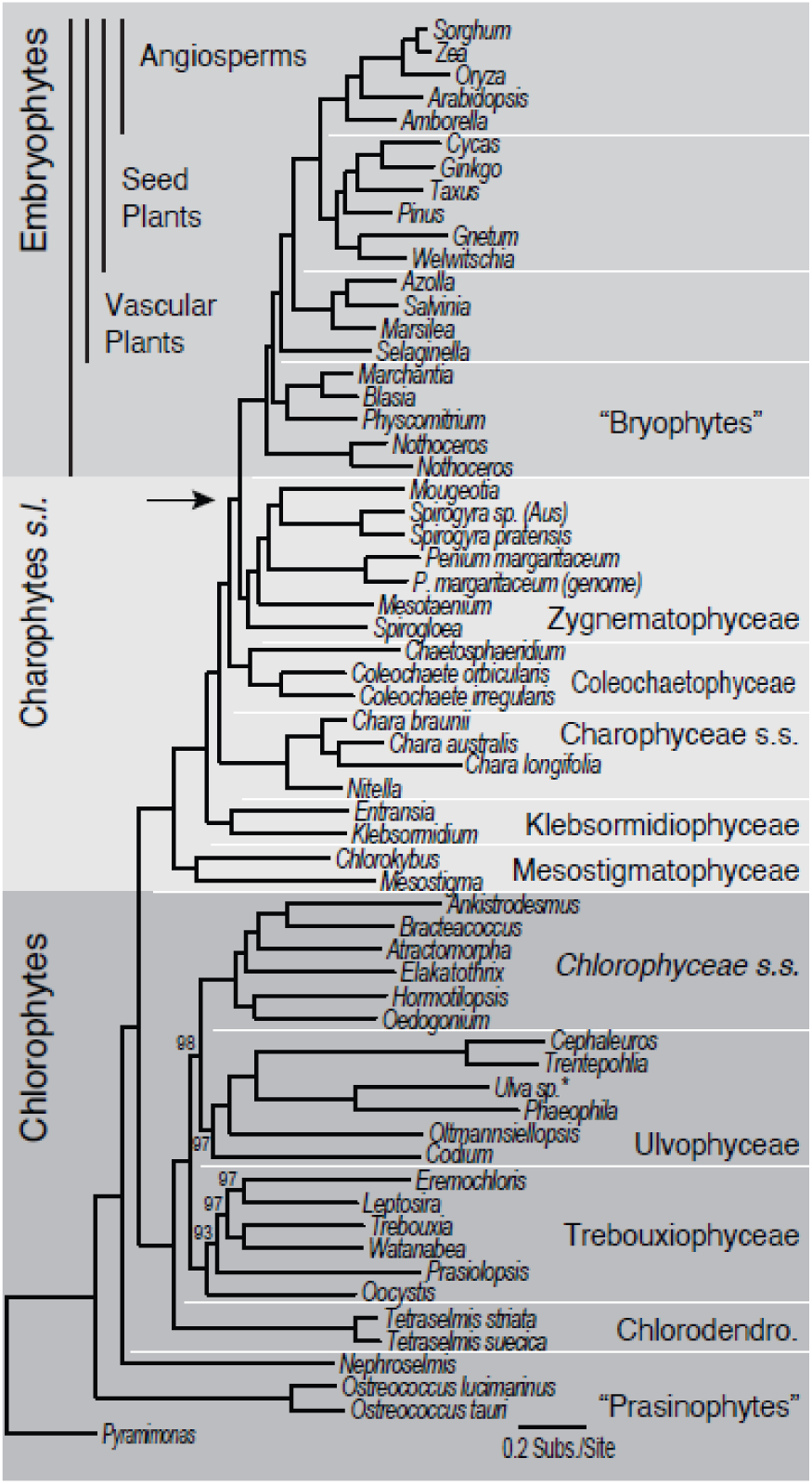
Phylogenetic Tree from the LG+F+R10 Analysis. Phylogenetic tree from the IQ-TREE LG+F+R10 analysis. The tree was rooted at *Pyramimonas parkeae*. Bootstrap support values less than 100 from ten thousand ultrafast bootstrap replicates are shown at each node.

**Figure 2:**
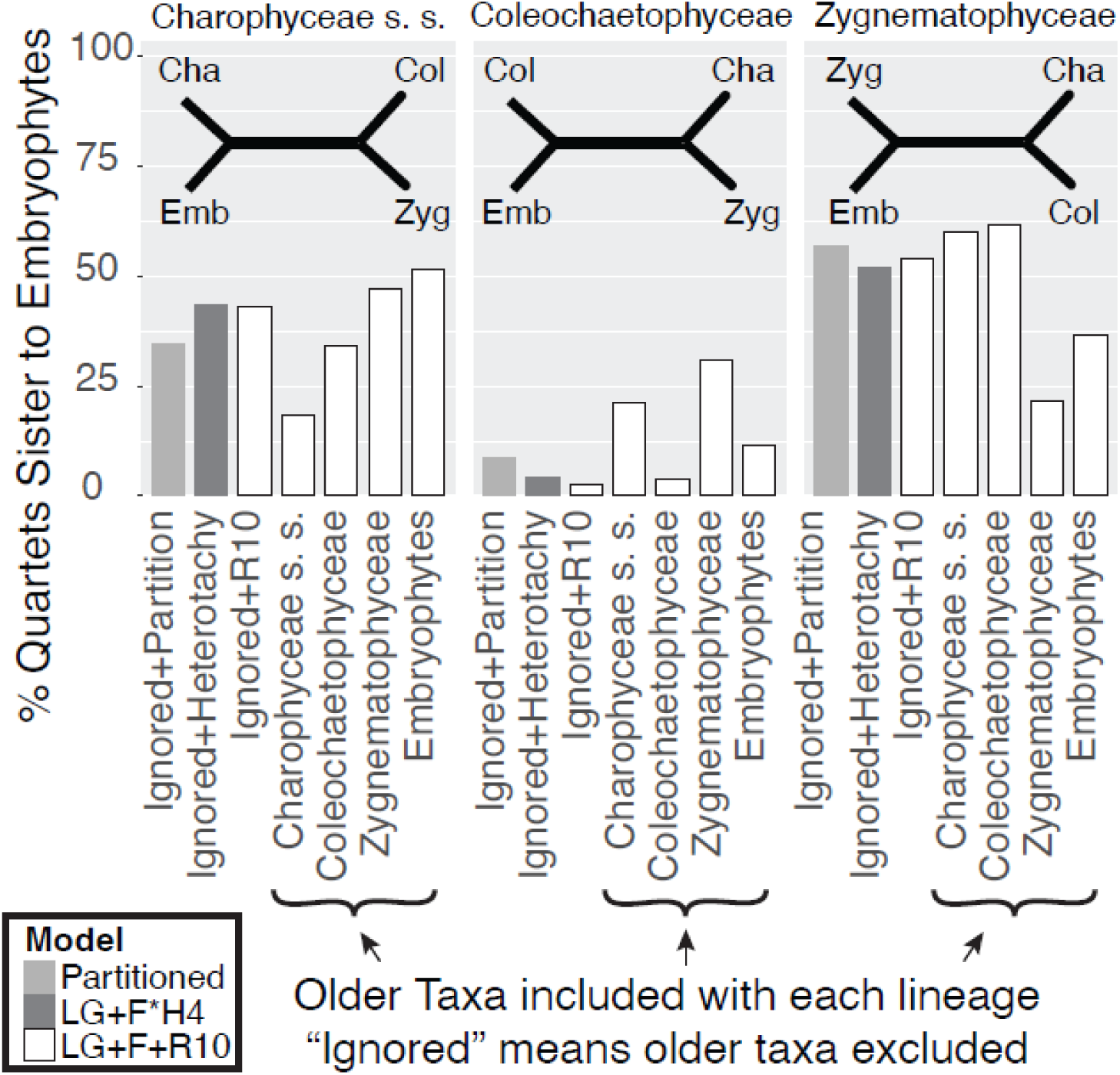
Likelihood Mapping Analysis. Percentage of likelihood quartets that support each of three charophyte lineages as the sister lineage to embryophytes. The corresponding four-taxon tree topology is shown at the top of each panel. The x axis describes where older taxa are included in the likelihood mapping analysis. The number of likelihood quartets for each different grouping of Older Taxa is as follows: Ignored = 1680, Charophyceae s. s. = 13440, Coleochaetophyceae = 17360, Zygnematophyceae = 8400, Embryophytes = 4032.

**Figure 3:**
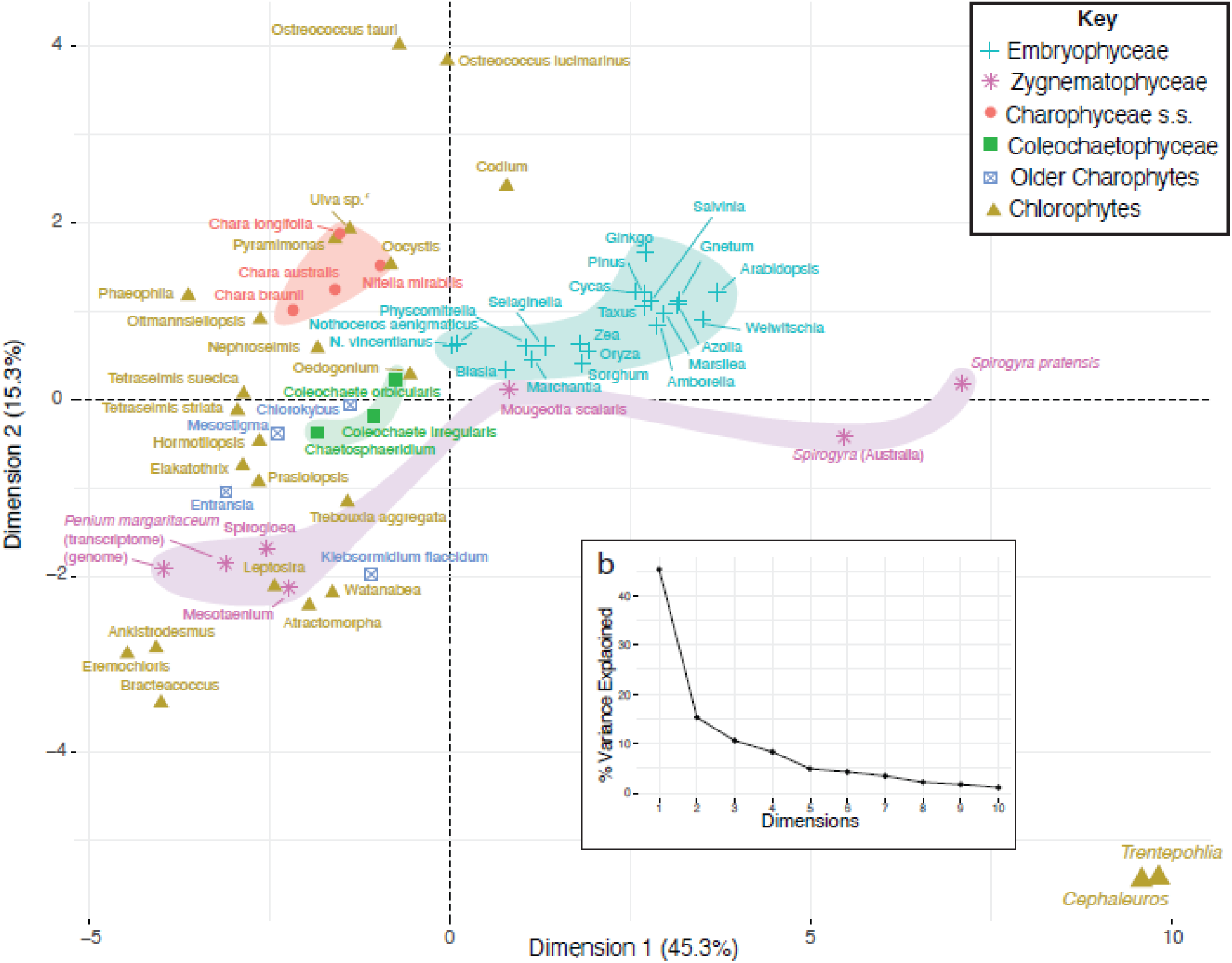
Principal Components Analysis of Amino Acid Frequencies in Orthologous Genes. Dimensions 1 and 2 of the principal components analysis of amino acid frequencies of orthologous genes included in the multiple sequence alignment. Inset scree plot shows the percentage of variation explained by each dimension.

### Phylogenetic analyses

The phylogenetic tree from the LG+F+R10 phylogenetic analysis is displayed in Figure 1. The likelihood analyses that ignored the older taxa resulted in a total of 1680 likelihood quartets. The percentages of likelihood quartets that support the Charophyceae s. s., Coleochaetophyceae, and Zygnematophyceae as the sister lineage to embryophytes are shown in Figure 2. Almost all of the phylogenetic analyses are generally in agreement on the overall topology of the tree, with only minor disagreements on specific branches. The nucleotide analysis with all codon positions shows the strongest departure from the other trees, especially in the Streptophyte lineages, where Coleochaetophyceae and Zygnematophyceae are non-monophyletic and *Selaginella* is shown as the outgroup to all other embryophyte lineages. These features are absent from the analysis done with only first and second codon positions. Of the three charophyte clades proposed thus far as sister to embryophytes, we recover strong and consistent support for Zygnematophyceae as the true sister lineage to land plants.

### Amino acid and codon composition analyses

The first dimension of all three principal components analyses explains around half of the variation in the data (45.3% for AA frequencies in taxa used for the trees; 49.3% for entire protein for all species; and 56.3% for codon frequencies), indicating that there is structure in the amino acid and codon frequencies among the Chloroplastida. In all three plots, the first principal component generally represents a gradual movement along the phylogeny from chlorophyte to charophyte to embryophyte. *Trentepohlia* and *Cephaleuros*, which use an alternate genetic nuclear code, cluster together far away from other chlorophyte taxa in these analyses.

### Protein domain and protein family searches

The ethylene signaling pathway protein domain searches are shown in Figure 4 and S14-S16. Among the algae, the CTR1 N-domain has the strongest signal in the Klebsormidiophyceae and Zygnematophyceae, the EIN2 signaling domain has the strongest signal in the Coleochaetophyceae and Zygnematophyceae, the EIN3 gene is equally strong in all charophyte lineages past the Mesostigmatophyceae, and the ETR1 ethylene binding domain has the strongest signal in the Zygnematophyceae. The NHX, CHX, KEA, and AAP gene family searches are shown in Figure 5 and S17-S19. The NHX family has a stronger signal in the charophytes than the chlorophytes and the CHX family is entirely absent from the chlorophytes. The KEA family appears to be universally present through the tree. The AAP family is absent from almost all of the algae, apart from two taxa in the Trebouxiophytes.

**Figure 4:**
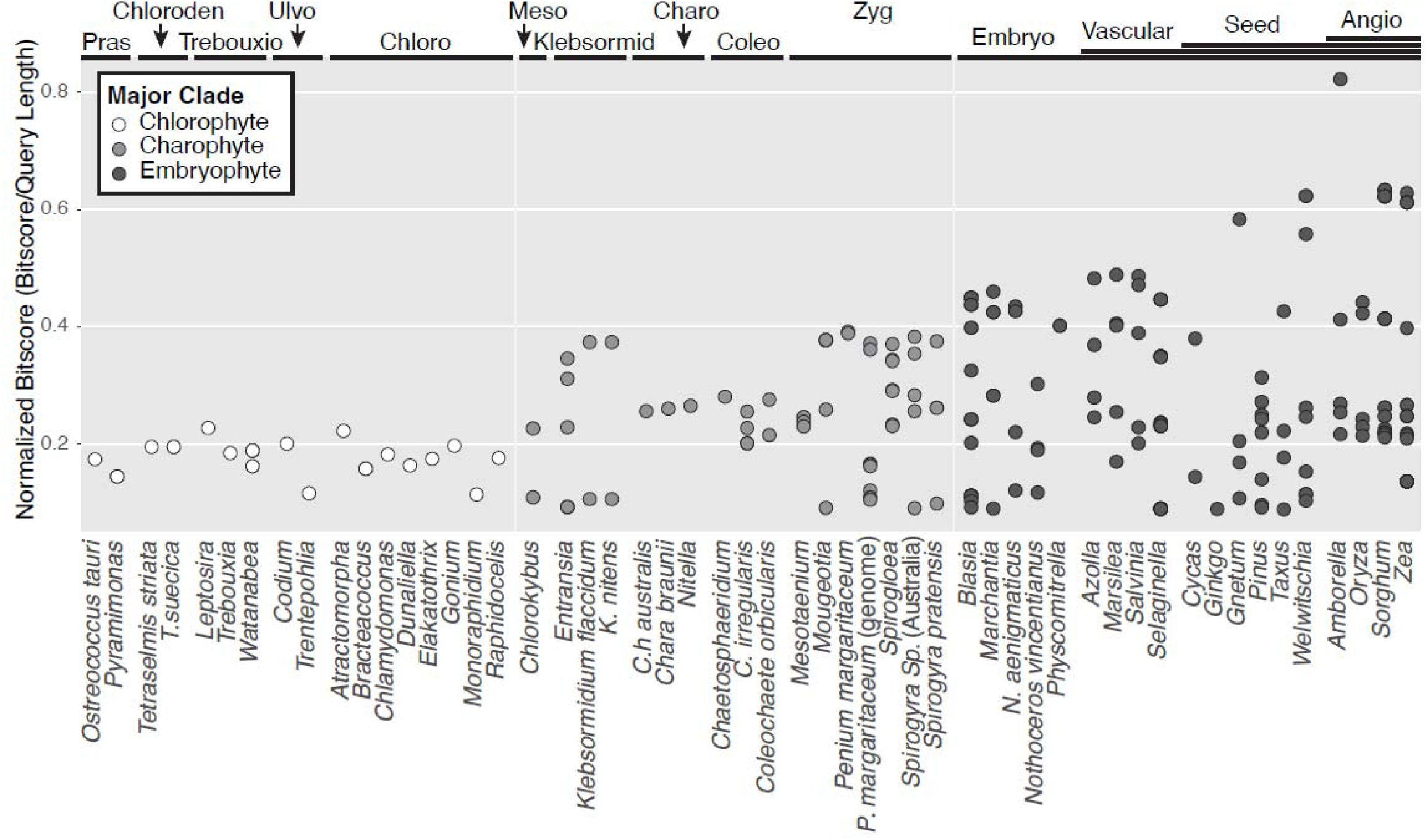
CTR1 N-terminal Domain Search. BLAST search of the dataset with the CTR1 N-terminal domain of the *Arabidopsis thaliana* sequence. Algal clade abbreviations going from left to right: Pras=Prasinophytes, Chloroden=Chlorodendrophyceae, Trebouxio=Trebouxiophyceae, Ulvo=Ulvophyceae, Chloro=Chlorophyceae, Meso=Mesostigmatophyceae, Klebsormid=Klebsormidiophyceae, Charo=Charophyceae, Coleochaeto=Coleochaetophyceae, Zygnemato=Zygnematophyceae. Embryophte taxa begin with the Bryophytes on the left and move across the tree towards the Angiosperms on the right.

**Figure 5:**
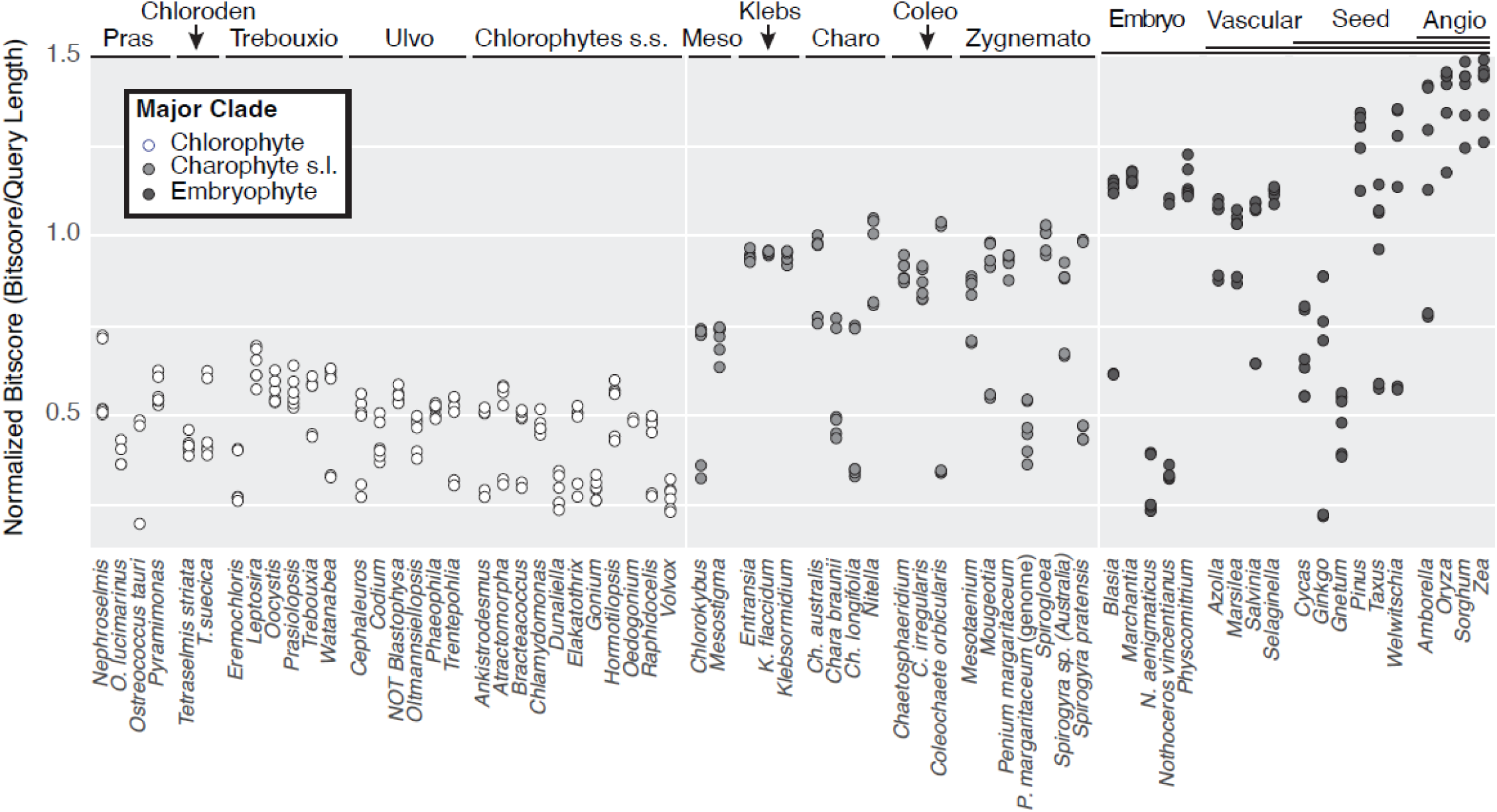
NHX Gene Family Searches. BLAST search of the dataset with the NHX gene family using *Arabidopsis thaliana* sequences (n=6). To reduce clutter, only the best hit for each individual gene in the family is retained for each taxa. Algal clade abbreviations going from left to right: Pras=Prasinophytes, Chloroden=Chlorodendrophyceae, Trebouxio=Trebouxiophyceae, Ulvo=Ulvophyceae, Chlorophytes s. s.=Chlorophyceae, Meso=Mesostigmatophyceae, Klebsormid=Klebsormidiophyceae, Charo=Charophyceae, Coleochaeto=Coleochaetophyceae, Zygnemato=Zygnematophyceae. Embryophte taxa begin with the Bryophytes on the left and move across the tree towards the Angiosperms on the right.

## Discussion

### Dataset quality and completeness

Since our phylogenetic analyses are reliant on a bioinformatic dataset drawn from many different sources, there will be differences between taxa originating in different extraction protocols, filtration cutoffs, and methods of assembly. Even within closely related species from our own *de novo* assemblies there are large differences, such as those between *Chaetosphaeridium globosum* and *Coleochaete orbicularis*. The *de novo* transcriptomes were built using RNA extracted from vegetative tissue and may have failed to include genes involved in specific biological functions.

### Green algae phylogenetics and the sister lineage to land plants

Our phylogenetic analyses found a green algal and broader Chloroplastida phylogeny that is consistent with previously published work by utilizing a dataset comprised of a large number of orthologous genes identified from both genomic and transcriptomic datasets, emphasizing the power of bioinformatic approaches in recovering deep evolutionary relationships (Cheon *et al*. 2018). Although recent multi-gene analyses show Ulvophyceae as being non-monophyletic, including placing *Codium fragile* as more closely related to the Chlorophyceae, we recover strong support for three monophyletic “core chlorophyte” clades in the IQ-TREE analyses (Trebouxiophyceae, Ulvophyceae, and Chlorophyceae) (Del Cortona *et al*. 2020; Fučíková *et al*. 2014). The ASTRAL analysis, however, places *Codium* as the outgroup to all the Chlorophyceae, a topology similar to that found by Li *et al*. 2021. The algal genomes used in HMM construction for ortholog identification all came from freshwater lineages (Chlorophyceae and various charophyte clades), which may have lessened their ability to detect genes suited for phylogenetic inference within the Ulvophyceae, many of which are green seaweeds and subaerial terrestrial organisms. Since we do not have as many Ulvophyceae taxa in our analysis as the Del Cortona paper, we are inclined to defer to their tree topology. We recommend additional focus on the Ulvophyceae and Chlorophyceae to parse those evolutionary relationships. Our analysis does corroborate the Del Cortona *et al*. 2020 and Li *et al*. 2021 placement of the Trebouxiophyceae as the outgroup to the other two “core chlorophyte” clades.

Our analyses generally found streptophytes to form a pectinate tree with Mesostigma and Chlorokybus forming a monophyletic group. Klebsormidiophyceae are the next major group in our analyses, but Glass *et al*. (2023) found *Interfilum* to form a clade branching before the Klebsormidiophyceae. The remaining lineages, sometimes called the “higher streptophytes”, i.e., Charophyceae s.s., Coleochaetophyceae, Zygnematophyceae, and embryophytes, generally branch in that order, with Zygnematophyceae being the sister taxon to embryophytes. However, we sometimes observed an alternate topology with Charophyceae s.s. appearing as the sister taxon to embryophytes, and zygnematophytes swapping places to below the coleochaetophytes. We only rarely observed other topologies, such as a monophyletic group consisting of Charophyceae s.s. and Coleochaetophyceae, which were ranked lower by the 15 5-taxon trees analysis. The two topologies where embryophytes group with either zygematophytes or charophytes s.s. were predominant.

### Insights into algal colonization of terrestrial environments

There is striking diversity of both amino acid and codon usage to be found in the Zygnematophyceae charophyte algae, with *Mougeotia* and the two *Spirogyra*, which are filamentous algae, clustering closer to embryophytes rather than with the remainder of the green algae, while the single-celled “desmids” (*Penium*, a true desmid, as well as “placoderm desmids” *Spiroglea* and *Mesotaenium)* had amino acid and codon usage more characteristic of the bulk of the green algae, both chlorophytes and charophytes. Strangely, codon usage in the group seems to follow morphology rather than phylogeny, although our sampling is not sufficient to make this ascertain with confidence. When all codon positions are maintained in the nucleotide phylogenetic analyses, *Mougeotia* and the two *Spirogyra* taxa remain as the sister lineage to embryophytes, whereas the other Zygnematophyceae are placed elsewhere in the tree. Given that filtering out third codon positions results in monophyletic Zygnematophyceae, we suspect that third codon positions may have reached mutational saturation, or perhaps have undergone selection for different nucleotide compositions, and thus are uninformative in our analysis of these deep evolutionary relationships. Additionally, *Nitella mirabilis* and the three *Chara* are closer to embryophytes on the first principal component than single-celled Zygnemtaophyceae and the older charophyte clades Mesostigmatophyceae and Klebsormidiophyceae. We advise caution in overinterpreting the any particular principal component from the three analyses due to the varying quality and completeness in the underlying data. Nevertheless, we hypothesize that Charophyceae s. s. and early embryophyte taxa have experienced convergent evolution towards a lifestyle in subaerial terrestrial environments, resulting in a more similar amino acid and codon usage profile. This may also explain the moderate support for the placement of the Charophyceae s. s. as the sister lineage to land plants in the likelihood mapping analyses. Furthermore, patterns of amino acid and codon usage provide important context for interpreting phylogenetic trees, as biases in codon composition are known to hinder accurate tree construction, as well as giving insight into the nature of large transcriptomic and genomic datasets (Cox *et al*. 2014).

Our ethylene pathway protein domain searches typically agree with previous work on the evolution of the ethylene signaling pathway (Ju *et al*. 2015). All of the domains (the CTR1 N-terminal domain, the EIN2 signaling domain, and the ETR1 ethylene binding domain), as well as the full sequence of the EIN3 gene, are absent in most of the chlorophyte lineages but homologs can be identified in various charophyte algae beginning with the Klebsormidiophyceae. These hits are particularly strong in the Zygnematophyceae. In the context of previous experimental manipulations of *Spirogyra,* it will be important to examine the function of these genes in diverse charophytes, because the functional ethylene perception pathway seems to have evolved during diversification of the streptophyte lineage. It is unclear what the role of this mechanism is in deeper-branching lineages. This observation also reinforces support for the hypothesis that the Zygnematophyceae are the sister lineage to land plants. The comparatively weak hits for the ETR1 ethylene-binding domain in the Charophyceae s. s. and Coleochaetophyceae may indicate loss of that particular gene and/or domain in those lineages or may mean the search is perceiving a different signal. Broader sampling in the Charophyceae and genomic resources for the Coleochaetophyceae will help resolve this question.

The four ion transporter gene families we examined show distinct patterns across the Chloroplastida phylogeny. First, the NHX gene family is broadly present in all clades, but charophyte sequences, as expected, show a greater degree of similarity to *Arabidopsis thaliana*. NHX genes are implicated in responses to salinity in various angiosperms (Akram *et al*. 2020; Fu *et al*. 2020; Yarra 2019). The homologous sequences in the charophyte algae may have a similar function or may have arisen in those lineages and been coopted by embryophytes for their current purposes (Phipps *et al*. 2021). The CHX gene family is present only in the charophyte algae, beginning with *Chlorokybus*. CHX genes are hypothesized to be involved with osmotic regulation and ion management during reproduction in angiosperms. It remains unknown, however, what function they might have in the different algae (Phipps *et al*. 2021 ;Sze *et al*. 2004). By contrast, the KEA gene family shares roughly the same degree of similarity to the *Arabidopsis thaliana* sequence across the entire phylogeny, up to and including the different prasinophyte taxa. KEA genes were thus present in a similar form to their current state in the ancestral Chloroplastida organism and may be a core part of stress responses to ion imbalances in all green organisms (Chen *et al*. 2015; Chanroj *et al*. 2012). Finally, the AAP searches show that the gene family is only present in embryophytes, although there is a puzzling signal in a branch of the Trebouxiophytes. Previous work done on AAP genes indicates that they are absent from the chlorophytes, although only *Volvox* and *Chlamydomonas* were examined (Tegeder and Ward 2012). These gene family searches are significant in that they represent a broad scanning of presence/absence of genes across the phylogeny. Ideally, they will pave the way for future laboratory manipulation in different algae, especially those not commonly studied.

### Conclusions

We have assembled and analyzed a dataset of over 60 species of Chloroplastida, including many algae species not commonly examined in the scientific literature. We successfully recover a phylogeny for these organisms and corroborate the recent placement of the Zygnematophyceae algae lineage as the sister lineage to land plants. We then show that amino acid and codon usage patterns of these species typically follow the structure of the phylogeny, although there are some notable exceptions such as the Trentepohliales. We conclude by scanning our dataset for homologous sequences to protein domains and gene families known to have important biological functions in land plants and show that those gene families are either present through the phylogeny or arose in the early charophyte algae.

## Acknowledgements

The authors would like to thank Endymion D. Cooper for collecting the RNA-seq data used to assemble the green algae transcriptomes, Caren Chang and Stephen M. Mount for helpful suggestions throughout the analysis, and Charles A. Goodman for insightful discussions on bioinformatics. We would also like to thank our collaborators for providing data: the Mary A. Bisson lab at the University of Buffalo, the Jaakko Hyvönen lab at the University of Helsinki, and the Fay-Wei Li lab at Cornell University. Special thanks to Thomas G. Doak and the National Center for Genome Analysis Support (NCGAS) at the University of Indiana for providing computational resources (National Science Foundation under Grant Nos. DBI-1062432 2011, ABI-1458641 2015, and ABI-1759906 2018 to Indiana University) and Heven Sze at the University of Maryland College Park for providing expertise on plant biology. This work was supported in part by NSF Grant # DEB-1036506 to CFD.

## Data Archive

RNA sequencing reads used in this project were deposited at the National Center for Biotechnology Information (NCBI) under accession PRJNA734590. Data files used in later analyses can be found at https://figshare.com/articles/dataset/Green_Algae_Phylogenetics_Data_Files/25230077

## Abbreviations

BUSCO: Benchmarking Universal Single-Copy Orthologs

