## Supplementary Information and Figures for "Investigating the Evolution of Green Algae with a Large Transcriptomic Dataset"

**Methods**

**Identification of *Ulva* from culture labeled *Blastophysa***

Although the species from the KU 295 culture was labeled *Blastophysa rhizopus* we identified it as an *Ulva* species via BLAST search. We searched the RubisCO sequence from our transcriptome assembly which returned a greater than 98 percent sequence identity against multiple *Ulva* species on NCBI. Supplementary figures still refer to the species as “*NOT Blastophysa rhizopus.”*

**Nucleotide Trees**

The corresponding coding nucleotide sequences for each of the amino acid sequences were used for additional phylogeny construction. Nucleotide coding sequences were aligned with MACSE v2.05 to ensure that first, second, and third codon positions were kept constant during the alignment process (Ranwez *et al.* 2011). Two separate nucleotide analyses were performed, one which retained all codon positions and the other which retained only first and second codon positions. As with the amino acid data, trimAl removed sites in the superalignment that contained a gap for 25% or more taxa and the ModelFinder option was employed in IQ-TREE to determine the optimal model (GTR+F+R10) for the two alignments.

**Gene Family Searches**

Specific domains in the search include the N-terminal domain of CTR1, the signaling domain of EIN2, and the ethylene-binding domain of ETR1. Since there is no distinguishing protein domain in the sequence coded for by EIN3, the entire sequence was searched. Gene family searches were performed for the sodium/hydrogen antiporter (NHX), cation/hydrogen exchanger (CHX), potassium efflux antiporter (KEA), and amino acid permease (AAP) gene families. 6 NHX genes, 4 CHX genes, 6 KEA genes, and 8 AAP genes were searched for a total of 24 genes from all families.

**Results**

**Phylogenetic Analysis**

Trees from the other IQTREE analyses, models used in the IQTREE partition analysis, and the ASTRAL gene tree reconciliation analysis are displayed in Figures S5-S8. The phylogenetic tree from the nucleotide analysis that included all codon positions is shown in Figure S9 and the phylogenetic tree from the nucleotide analysis that included only first and second codon positions is shown in Figure S10. Bootstrap support is shown at each node in each of the trees constructed in IQ-TREE; nodes in the ASTRAL tree show local posterior probability support. The ranking of the 15 possible five-taxon trees is shown in Figure S11.

**Tables**

**Table S1: Algae species assembled *de novo***

| **Species** | **Major Clade** | **Minor Clade** | **Media** | **Temperature ^o^C** | **Collection** | **NCBI Accession** | **Contigs** |
| --- | --- | --- | --- | --- | --- | --- | --- |
| *Ankistrodesmus falcatus* | Chlorophyte | Chlorophyceae | GWH | 18 | UTEX101 | SRX13042245 | 52673 |
| *Atractomorpha echinata* | Chlorophyte | Chlorophyceae | GWH | 18 | UTEX LB2309 | SRX13042246 | 29637 |
| *Bracteacoccus aerius* | Chlorophyte | Chlorophyceae | GWH | 18 | UTEX1250 | SRX13042257 | 68993 |
| *Cephaleuros parasiticus* | Chlorophyte | Ulvophyceae | Solid GWH | 18 | SAG 73.90 | SRX13042268 | 40824 |
| *Chaetospharidium globosum* | Charophyte | Coleochaetophyceae | DY III | 18 | SAG 26.98 | SRX13042271 | 21568 |
| *Coleochaete orbicularis* | Charophyte | Coleochaetophyceae | GWH | 18 | LB422 | SRX13042272 | 34658 |
| *Elakatothrix viridis* | Chlorophyte | Chlorophyceae | GWH | 18 | SAG 9.94 | SRX13042273 | 25019 |
| *Entransia* | Charophyte | Klebsormidiophyceae | DY III | 18 | Delwiche Lab | SRX13042274 | 36135 |
| *Eremochloris sphaerica* | Chlorophyte | Trebouxiophyceae | GWH | 18 | Delwiche Lab | SRX13042275 | 34165 |
| *Hormotilopsis gelatinosa* | Chlorophyte | Chlorophyceae | GWH | 18 | UTEX B104 | SRX13042276 | 48998 |
| *Klebsormidium flaccidum* | Charophyte | Klebsormidiophyceae | BBM | 18 | UTEX 321 | SRX13042247 | 52970 |
| *Leptosira* | Chlorophyte | Trebouxiophyceae | GWH | 18 | UTEX 333 | SRX13042248 | 43769 |
| *Mesostigma viride* | Charophyte | [Mesostigmatophyceae](https://en.wikipedia.org/wiki/Mesostigmatophyceae) | GWH | 18 | NIES 995 | SRX13042249 | 27491 |
| *Mougeotia scalaris* | Charophyte | Zygnematophyceae | GWH | 18 | SAG 164.90 | SRX13042250 | 29610 |
| *Nephroselmis pyriformis* | Chlorophyte | Prasinophyta | 32 ppt f/2 | 16 | CCMP 717 | SRX13042251 | 22703 |
| *Nitella mirabilis lower* | Charophyte | Charophyceae | N/A | N/A | N/A | SRX13042252 | 39680 |
| *Nitella_mirabilis upper* | Charophyte | Charophyceae | N/A | N/A | N/A | SRX13042253 | 39680 |
| *Ulva sp. (culture was labeled Blastophysa)* | Chlorophyte | Ulvophyceae | N/A | 10 | KU 295 | SRX13042254 | 20570 |
| *Oedogonoium cardiacum* | Chlorophyte | Chlorophyceae | GWH | 18 | UTEX LB40 | SRX13042255 | 26857 |
| *Oltmannsiellopsis unicellularis* | Chlorophyte | Ulvophyceae | 30 ppt L1 | 16 | SCCAP K-2050 | SRX13042256 | 27443 |
| *Oocystis solitaria* | Chlorophyte | Trebouxiophyceae | GWH | 18 | SAH 83.80 | SRX13042258 | 49739 |
| *Penium margaritaceum* | Charophyte | Zygnematophyceae | GWH | 18 | SAG 22.82 | SRX13042259 | 46166 |
| *Phaeophila dendroides* | Chlorophyte | Ulvophyceae | 32 ppt L1 | 18 | CCMP 2372 | SRX13042260 | 25287 |
| *Prasiolopsis* | Chlorophyte | Trebouxiophyceae | Solid GWH | 18 | SAG 84-81 | SRX13042261 | 37064 |
| *Pyramimonas parkeae* | Chlorophyte | Prasinophyta | 32 ppt f/2 | 16 | CCMP 726 | SRX13042262 | 24787 |
| *Spirogyra pratensis* | Charophyte | Zygnematophyceae | GWH | 18 | UTEX 921 | SRX13042263 | 13470 |
| *Spirogyra Au1* | Charophyte | Zygnematophyceae | GWH | 18 | Delwiche Lab | SRX13042264 | 29533 |
| *Tetraselmis striata* | Chlorophyte | Chlorodendrophyceae | 32 ppt f/2 | 18 | SAG 41.85 | SRX13042265 | 37356 |
| *Tetraselmis suecica* | Chlorophyte | Chlorodendrophyceae | 32 ppt f/2 | 18 | PLY 305 | SRX13042266 | 30017 |
| *Trebouxia aggregata* | Chlorophyte | Trebouxiophyceae | GWH | 18 | SAG 219-1d | SRX13042267 | 62197 |
| *Trentepohlia annulata* | Chlorophyte | Ulvophyceae | Solid GWH | 18 | SAG 20.94 | SRX13042269 | 38824 |
| *Watanabea reniformis* | Chlorophyte | Trebouxiophyceae | GWH | 18 | SAG 211-9b | SRX13042270 | 70041 |

**Table S2: Additional taxa included in the phylogenetic analysis**

| **Species** | **Major Clade** | **Minor Clade** | **Data Type** | **Data Source** | **Contigs** |
| --- | --- | --- | --- | --- | --- |
| *Amborella trichopoda* | Embryophyte | Angiosperm | Genome | NCBI | 32950 |
| *Arabidopsis thaliana* | Embryophyte | Angiosperm | Genome | NCBI | 63286 |
| *Azolla filiculoides* | Embryophyte | Angiosperm | Genome | Fernbase | 30030 |
| *Blasia* | Embryophyte | Pre-Vascular | Transcriptome | Collaborator | 89464 |
| *Chara australis* | Charophyte | Charophyceae | Transcriptome | Collaborator | 31181 |
| *Chara braunii* | Charophyte | Charophyceae | Genome | NCBI | 109568 |
| *Chara longifolia* | Charophyte | Charophyceae | Transcriptome | Collaborator | 25167 |
| *Chlorokybus atmophyticus* | Charophyte | Chlorokybophyceae | Transcriptome | OneKP | 28131 |
| *Codium fragile* | Chlorophyte | Ulvophyceae | Transcriptome | OneKP | 20311 |
| *Coleochaete irregularis* | Charophyte | Coleochaetophyceae | Transcriptome | OneKP | 40991 |
| *Cycas micholitzii* | Embryophyte | Gymnosperm | Transcriptome | OneKP | 20176 |
| *Ginkgo biloba* | Embryophyte | Gymnosperm | Transcriptome | OneKP | 22056 |
| *Gnetum montanum* | Embryophyte | Gymnosperm | Transcriptome | OneKP | 25386 |
| *Marchantia polymorpha* | Embryophyte | Pre-Vascular | Genome | Genome Website | 43273 |
| *Marsilea* | Embryophyte | Early Vascular | Transcriptome | Collaborator | 51533 |
| *Mesotaenium endlicherianum* | Charophyte | Zygnematophyceae | Genome | Genome Website | 33631 |
| *Nothoceros aenigmaticus* | Embryophyte | Pre-Vascular | Transcriptome | OneKP | 33555 |
| *Nothoceros vincentianus* | Embryophyte | Pre-Vascular | Transcriptome | OneKP | 35360 |
| *Oryza sativa* | Embryophyte | Angiosperm | Genome | NCBI | 64394 |
| *Ostreococcus lucimarinus* | Chlorophyte | Prasinophyta | Genome | NCBI | 19593 |
| *Ostreococcus tauri* | Chlorophyte | Prasinophyta | Genome | NCBI | 20823 |
| *Penium margaritaceum* | Charophyte | Zygnematophyceae | Genome | Genome Website | 131645 |
| *Physcomitrium patens* | Embryophyte | Pre-Vascular | Genome | NCBI | 84741 |
| *Pinus taeda* | Embryophyte | Gymnosperm | Transcriptome | OneKP | 48170 |
| *Salvinia cucullata* | Embryophyte | Early Vascular | Transcriptome | Fernbase | 37131 |
| *Selaginella moellendorffii* | Embryophyte | Early Vascular | Genome | NCBI | 107591 |
| *Sorghum bicolor* | Embryophyte | Angiosperm | Genome | NCBI | 104545 |
| *Spiroglea muscilosa* | Charophyte | Zygnematophyceae | Genome | Genome Website | 83089 |
| *Taxus baccata* | Embryophyte | Gymnosperm | Transcriptome | OneKP | 25467 |
| *Welwitschia mirabilis* | Embryophyte | Gymnosperm | Transcriptome | OneKP | 28839 |
| *Zea mays* | Embryophyte | Angiosperm | Genome | NCBI | 121429 |

**Table S3: Algal genomes used in hidden Markov model construction for ortholog identification**

| **Species** | **Major Clade** | **Minor Clade** | **Data Source** | **Contigs** |
| --- | --- | --- | --- | --- |
| *Chara braunii* | Charophyte | Charophyceae | NCBI | 109568 |
| *Chlamydomonas reinhardtii* | Chlorophyte | Chlorophyceae | NCBI | 93864 |
| *Dunaliella salina* | Chlorophyte | Chlorophyceae | NCBI | 49998 |
| *Gonium pectorale* | Chlorophyte | Chlorophyceae | NCBI | 63197 |
| *Klebsormidium nitens* | Charophyte | Klebsormidiophyceae | NCBI | 47449 |
| *Monoraphidium neglectum* | Chlorophyte | Chlorophyceae | NCBI | 52254 |
| *Penium margaritaceum* | Charophyte | Zygnematophyceae | Genome Website | 131645 |
| *Raphidocelis subcapitata* | Chlorophyte | Chlorophyceae | NCBI | 61044 |
| *Volvox carteri* | Chlorophyte | Chlorophyceae | NCBI | 49552 |

**Table S4:**

| **Grouping** | **Taxa** |
| --- | --- |
| Charophyceae s. s. | *Chara australis, Chara braunii, Chara longifolia, Nitella mirabilis* |
| Coleochaetophyceae | *Chaetosphaeridium globosum, Coleochaete irregularis, Coleochaete orbicularis* |
| Embryophytes | *Amborella trichopoda, Arabidopsis thaliana, Azolla filiculoides, Blasia, Cycas micholitzii, Ginkgo biloba, Gnetum montanum, Marchantia polymorpha, Marsilea, Nothoceros aenigmaticus, Nothoceros vincentianus, Oryza sativa, Physcomitrium patens, Pinus taeda, Salvinia cucullata, Selaginella moellendorffii, Sorghum bicolor, Taxus baccata, Welwitschia mirabilis, Zea mays* |
| Older Taxa | *Ankistrodesmus falcatus, Atractomorpha echinata, Bracteacoccus aerius, Cephaleuros parasiticus, Chlorokybus atmophyticus ,Codium fragile, Elakatothrix viridis, Entransia Eremochloris sphaerica, Hormotilopsis gelatinosa, Klebsormidium flaccidum, Leptosira, Mesostigma viride, Ulva sp. (from a mislabeled culture, labeled “Not Blastophysa” in some legacy files and analyses), Nephroselmis pyriformis, Oedogonium cardiacum, Oltmannsiellopsis unicellularis, Oocystis solitaria, Ostreococcus lucimarinus, Ostreococcus tauri Phaeophila dendroides, Prasiolopsis, Pyramimonas parkeae, Tetraselmis striata, Tetraselmis suecica, Trebouxia aggregata, Trentepohlia annulata, Watanabea reniformis* |
| Zygnematophyceae | *Mesotaenium endlicherianum, Mougeotia scalaris, Penium margaritaceum genome, Penium margaritaceum, Spirogloea muscicola, Spirogyra Aus, Spirogyra pratensis* |

**Supplementary Figures**

**Supplementary Figure 1: Orthogroup Presence Across Taxa**

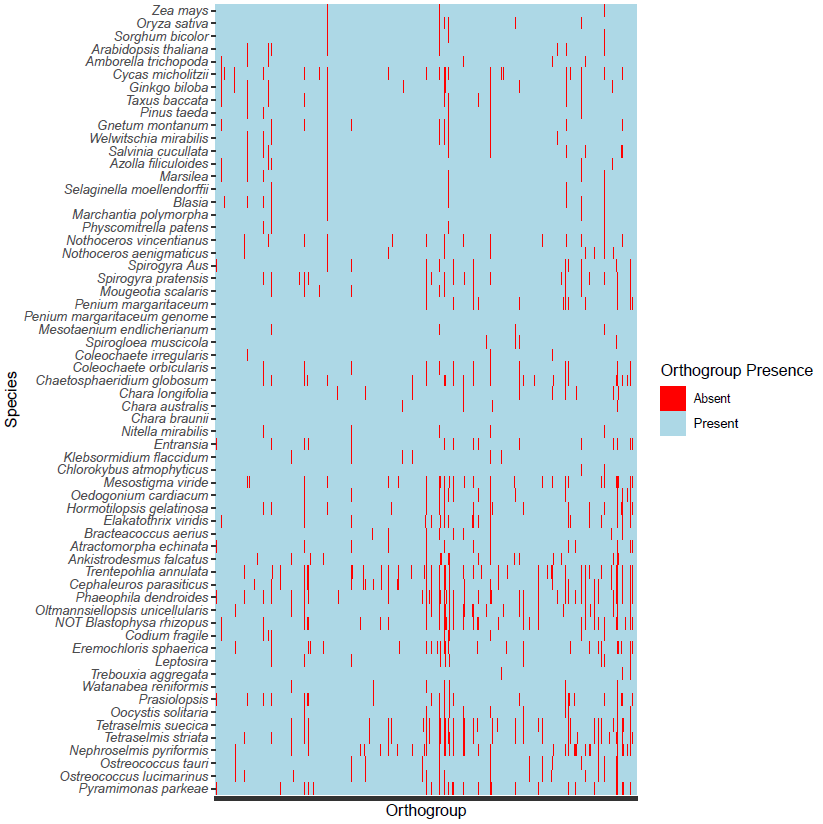

Supplementary Figure 1: Orthogroup presence and absence across taxa in the phylogenetic dataset. The *Penium margaritaceum* and *Chara braunii* genomes were used in both HMM construction and in the later phylogenetic analysis.

**Supplementary Figure 2: BUSCO Scores of Taxa in the Phylogenetic Analysis**

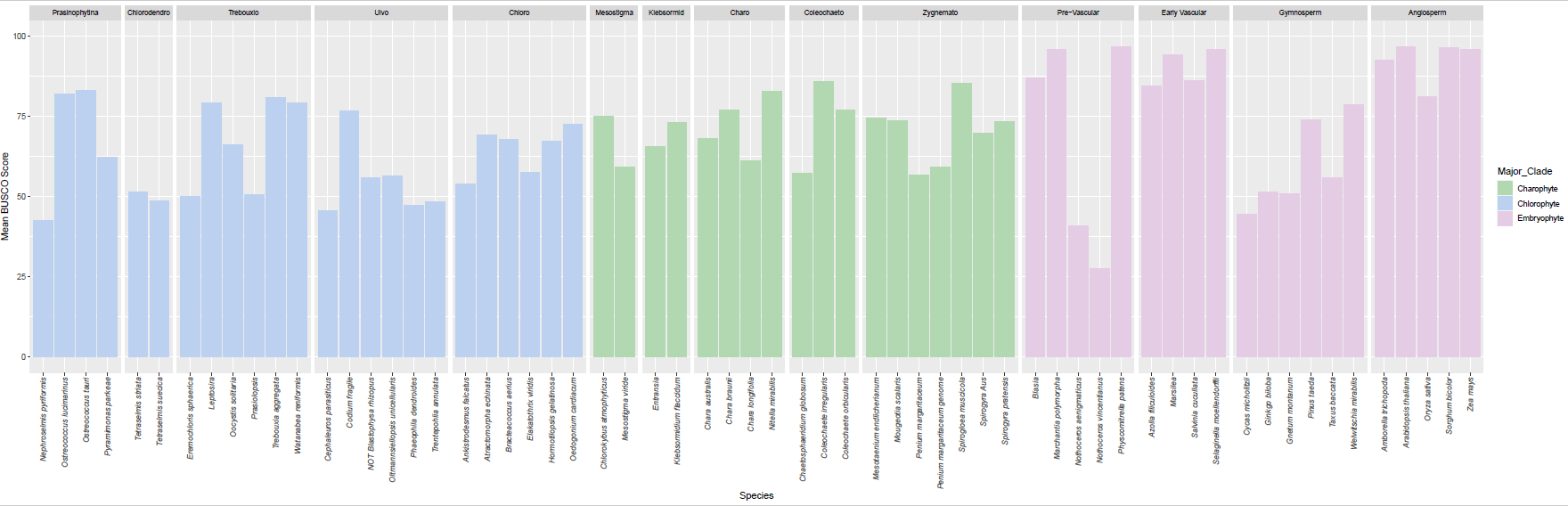

Supplementary Figure 2: Mean BUSCO scores across the Viridiplantae, Chlorophyte, and Eukaryote BUSCO databases for each taxa in the phylogenetic dataset. Clade abbreviations going from left to right: Chlorodendro=Chlorodendrophyceae, Trebouxio=Trebouxiophyceae, Ulvo=Ulvophyceae, Chloro=Chlorophyceae, Mesostigma=Mesostigmatophyceae, Klebsormid=Klebsormidiophyceae, Charo=Charophyceae, Coleochaeto=Coleochaetophyceae, Zygnemato=Zygnematophyceae.

**Supplementary Figure 3: BUSCO Scores for Algal Genomes**

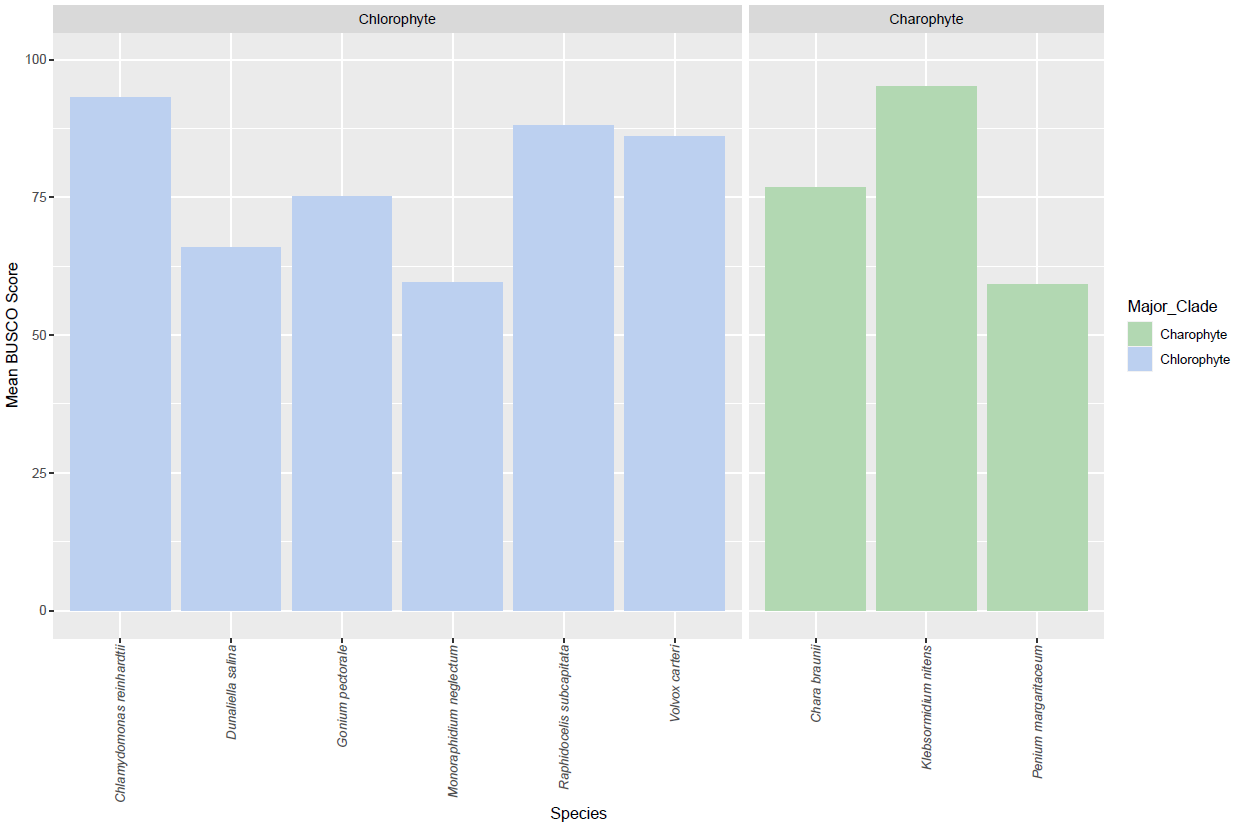

Supplementary Figure 3: Mean BUSCO scores across the Viridiplantae, Chlorophyte, and Eukaryote BUSCO databases for each algal taxa used for orthogroup identificaiton and hidden Markov model construction.

**Supplementary Figure 4: Percentage of Gaps in the SuperAlignment**

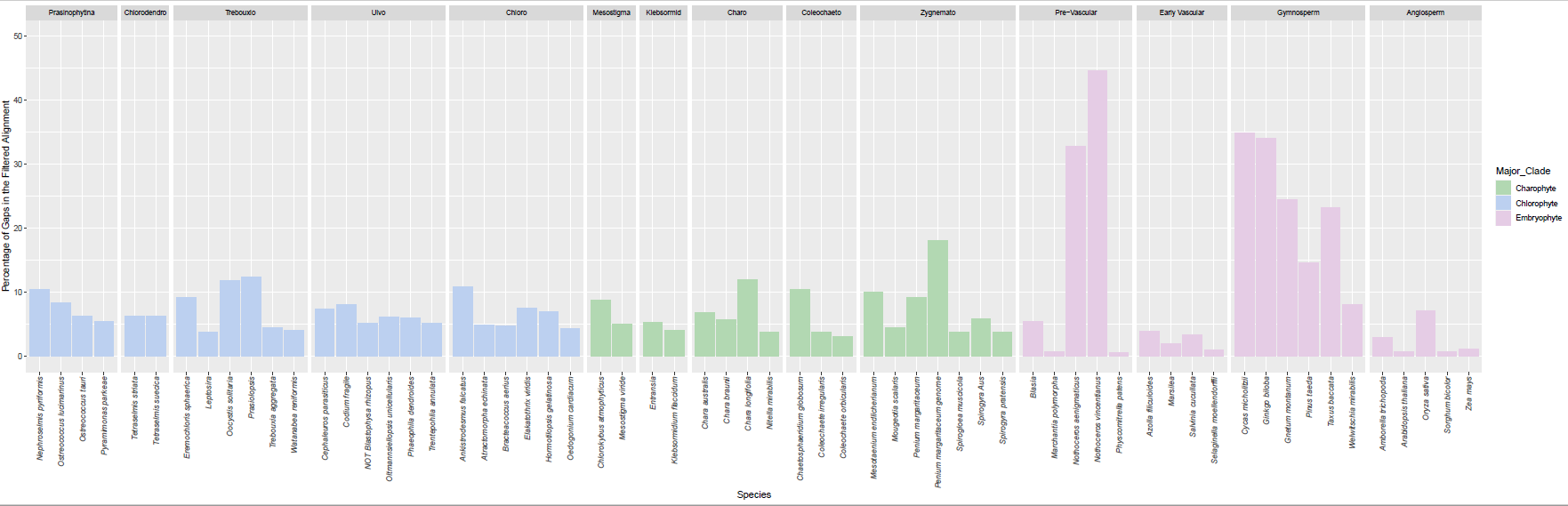

Supplementary Figure 4: Percentage of gaps in each taxa in the phylogenetic analysis after trimAl filtering.

**Supplementary Figure 5: Phylogenetic Tree from the LG+F*H4 Analysis**

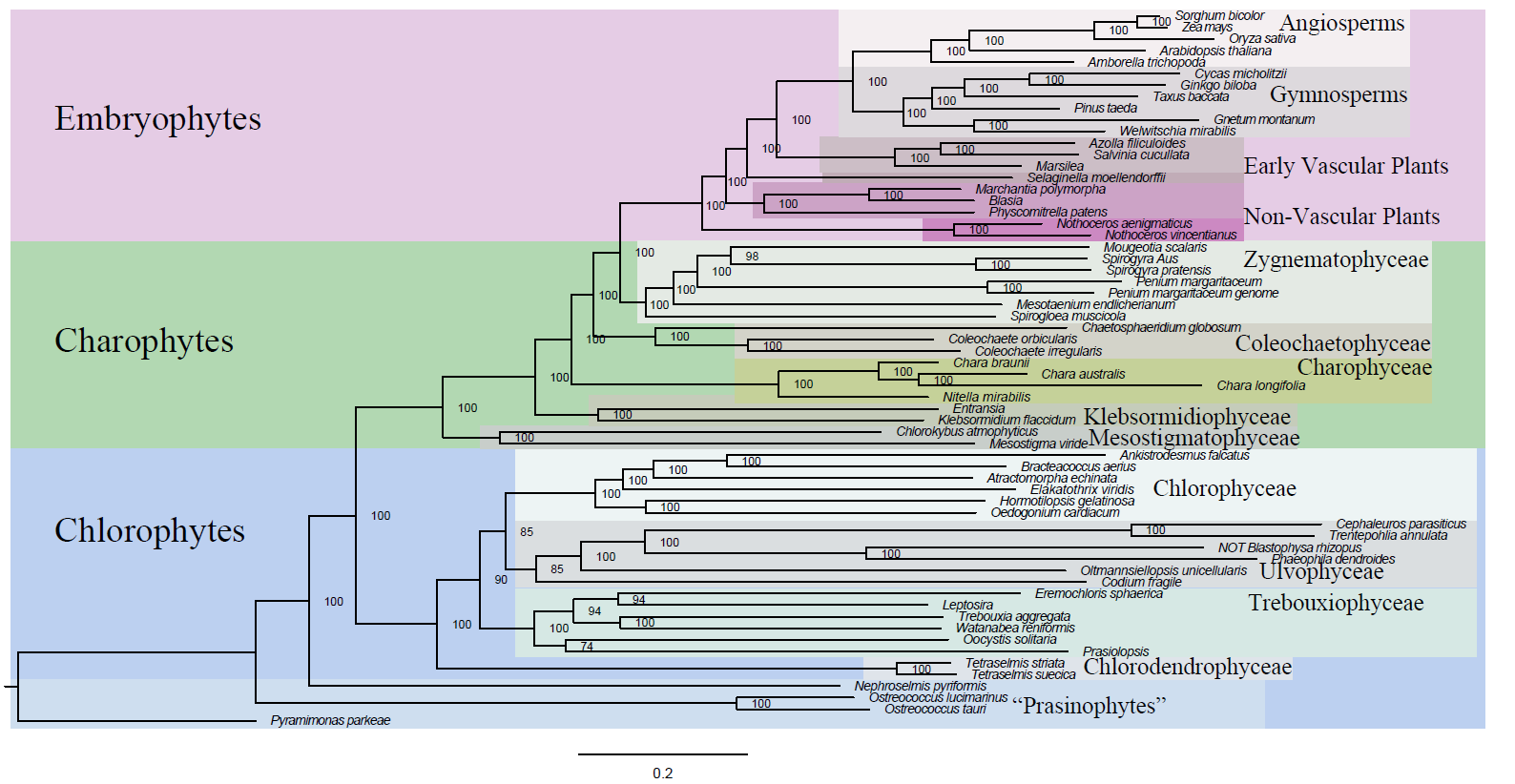

Supplementary Figure 5: Phylogenetic tree from the IQ-TREE LG+F*H4 analysis. The tree was rooted at *Pyramimonas parkeae*. Bootstrap support values from ten thousand ultrafast bootstrap replicates are shown at each node.

**Supplementary Figure 6: Distribution of Optimal Models for each individual Gene Tree**

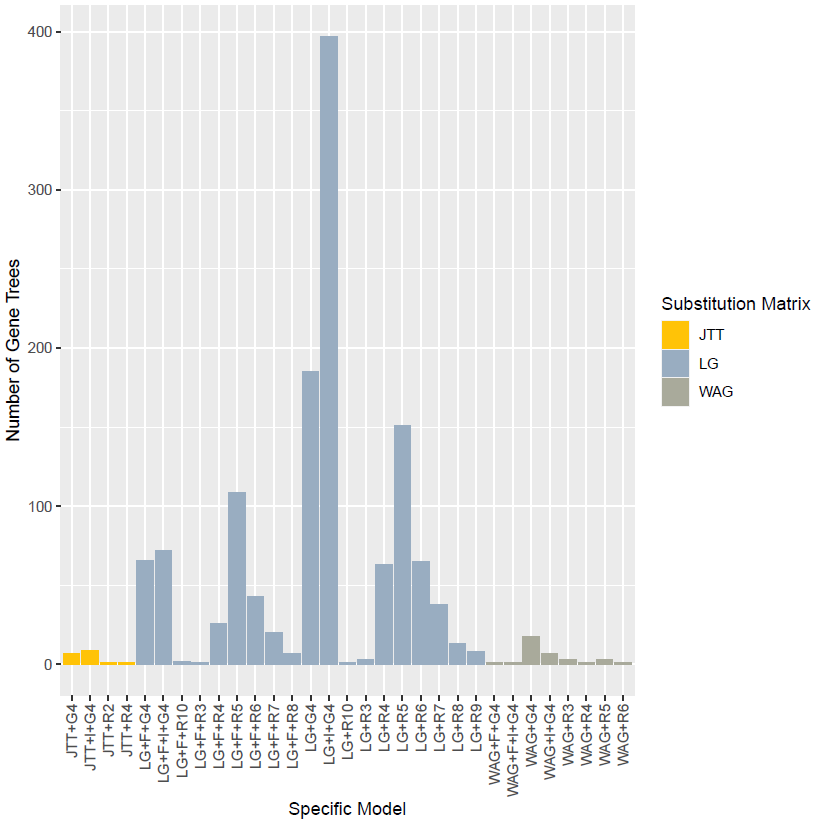

Supplementary Figure 6: Distribution of gene models found for each of the 1323 gene trees. Gene trees are organized by the amino acid substitution matrix (JTT, LG, or WAG) and number and type of rate heterogeneity parameters. I indicates that the model uses a proportion of invariant sites, F indicates that the model uses empirical amino acid frequencies, G indicates that the rate parameters are drawn from a gamma distribution, and R indicates that the distribution of rate parameters is nonparametric.

**Supplementary Figure 7: Phylogenetic Tree from the Partitioned Analysis**

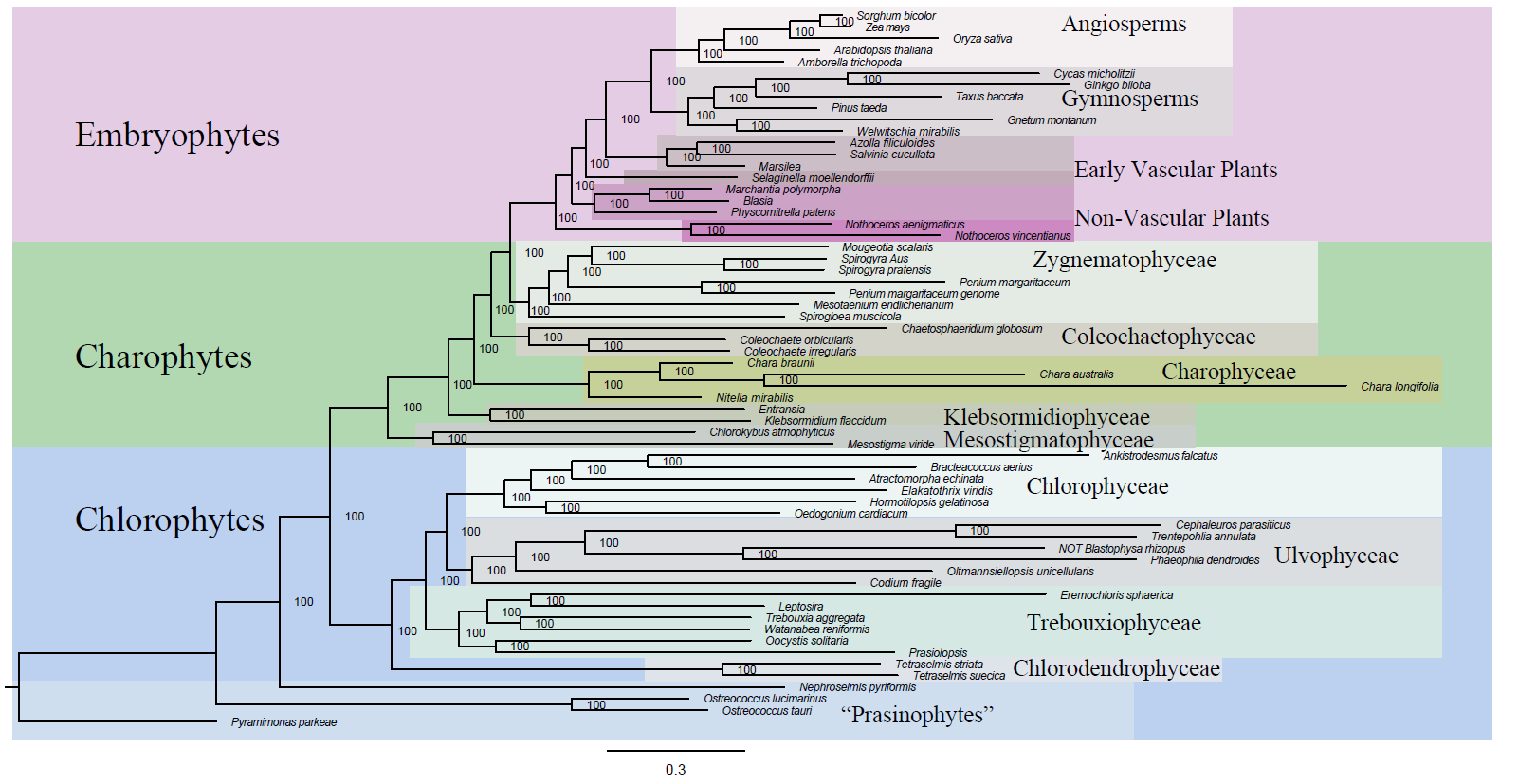

Supplementary Figure 7: Phylogenetic tree from the IQ-TREE partitioned analysis. The tree was rooted at *Pyramimonas parkeae*. Bootstrap support values from ten thousand ultrafast bootstrap replicates are shown at each node.

**Supplementary Figure 8: ASTRAL Phylogenetic Tree**

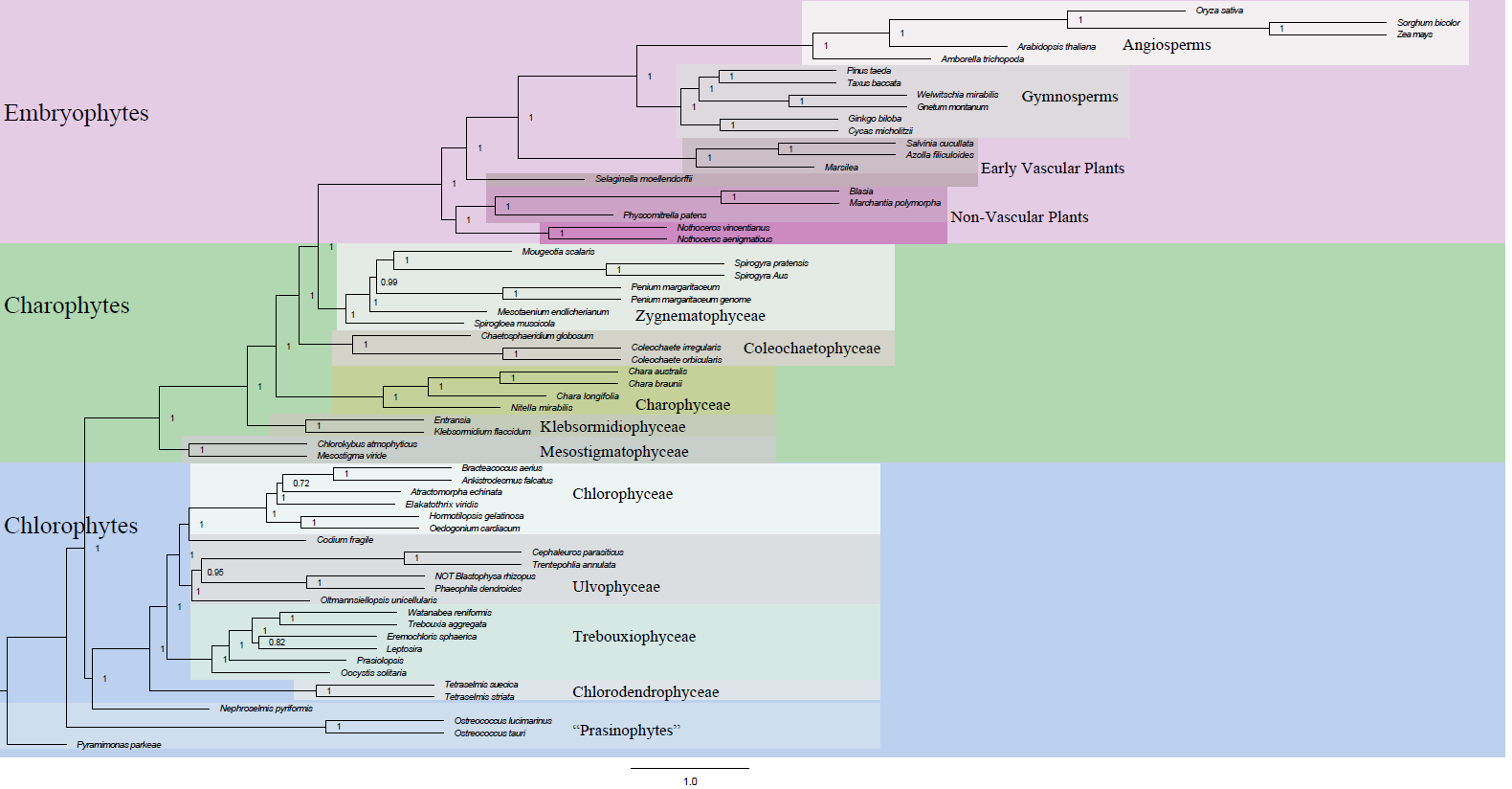

Supplementary Figure 8: ASTRAL phylogenetic tree constructed with 1323 individual gene trees. The tree was rooted at *Pyramimonas parkeae*. Node labels show the local posterior probability for that branch.

**Supplementary Figure 9: Nucleotide Tree with All Codon Positions**

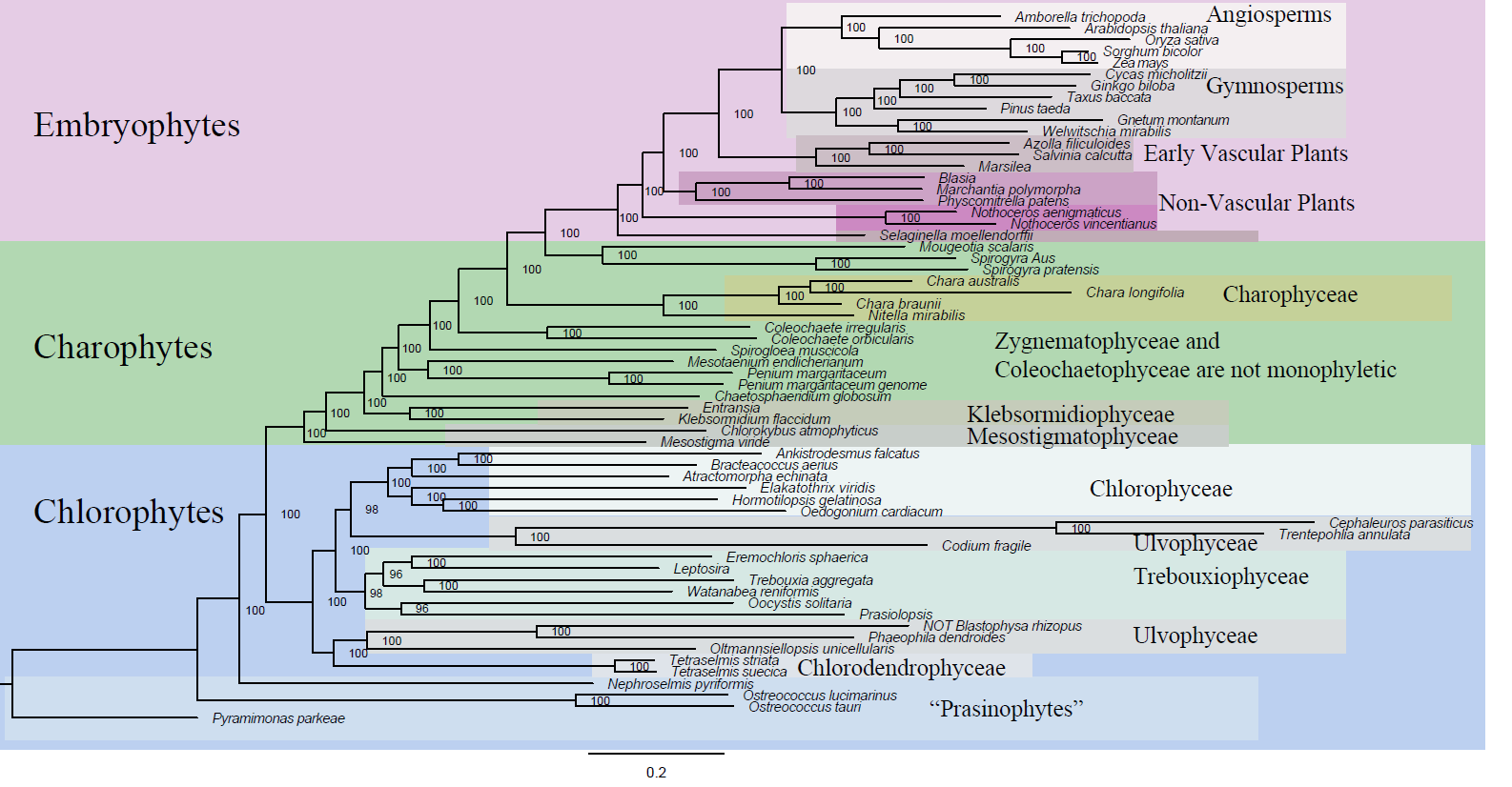

Supplementary Figure 9: Phylogenetic tree from the IQ-TREE GTR+F+R10 nucleotide analysis with all codon positions. The tree was rooted at *Pyramimonas parkeae*. Bootstrap support values from ten thousand ultrafast bootstrap replicates are shown at each node.

**Supplementary Figure 10: Nucleotide Tree with First and Second Codon Positions**

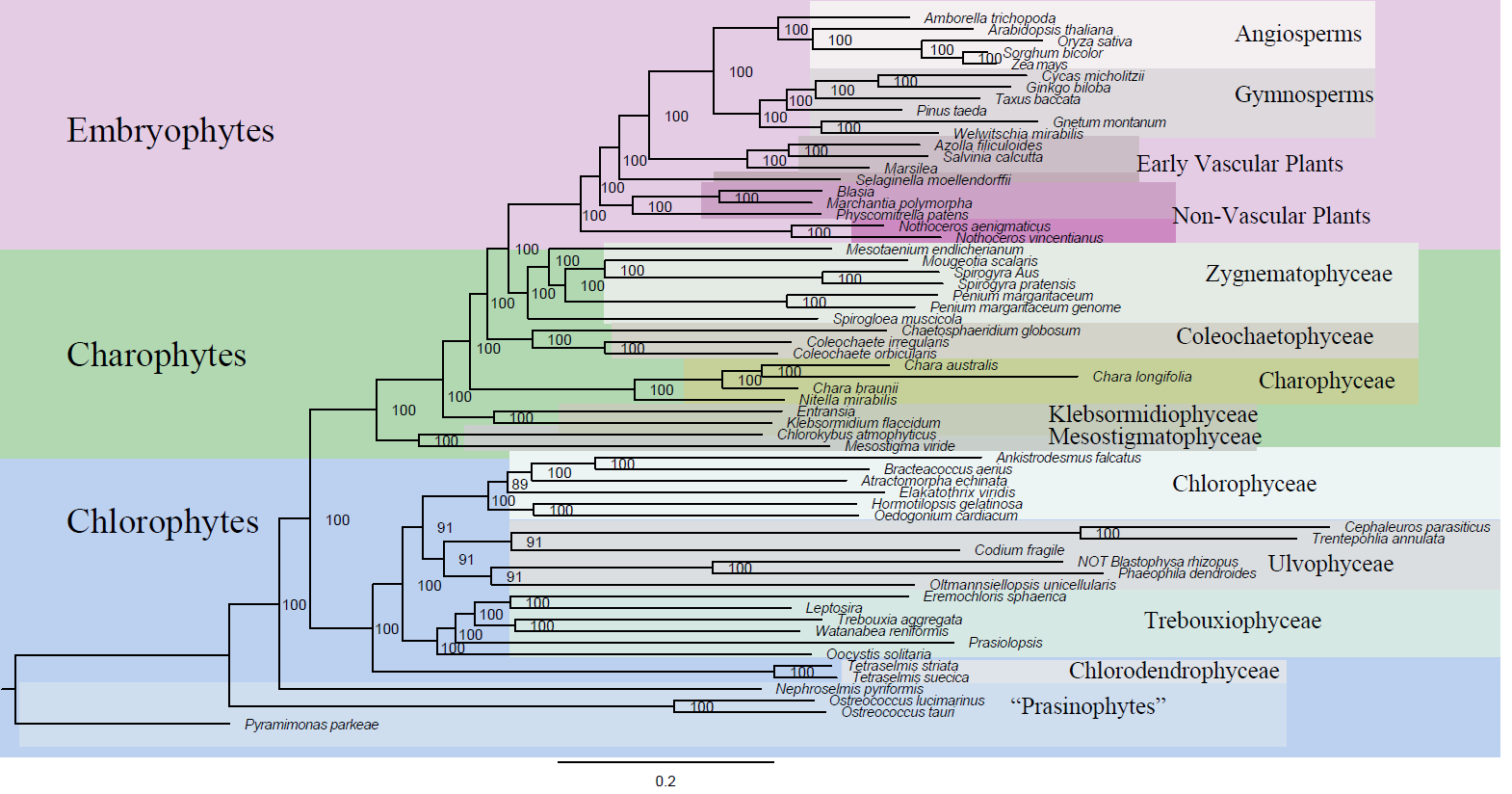

Supplementary Figure 10: Phylogenetic tree from the IQ-TREE GTR+F+R10 nucleotide analysis with first and second codon positions. The tree was rooted at *Pyramimonas parkeae*. Bootstrap support values from ten thousand ultrafast bootstrap replicates are shown at each node.

**Supplementary Figure 11: Analysis of 15 Possible Five-Taxon Trees**

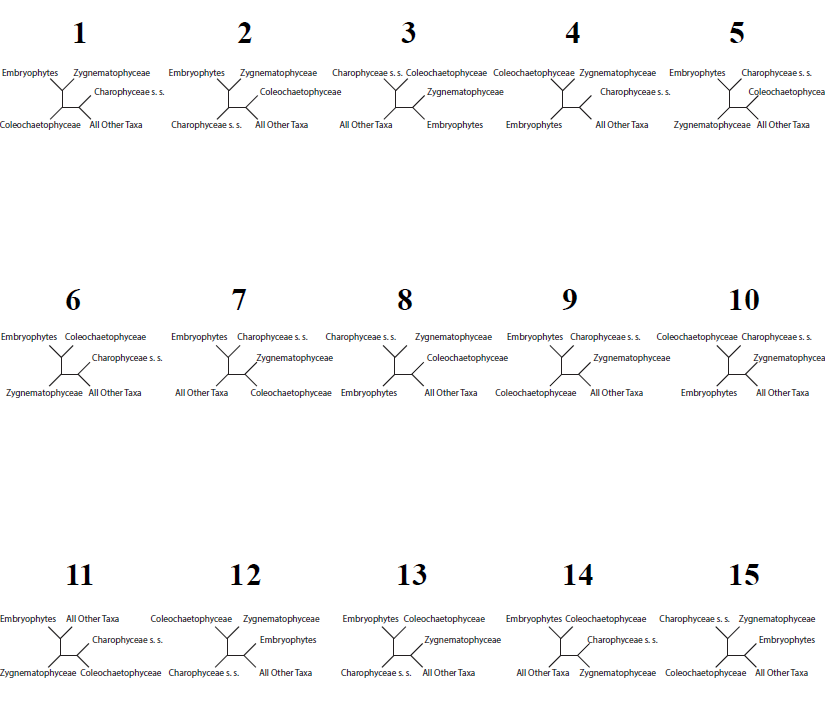

Supplementary Figure 11: Ranking, according to log-likelihood, of each of the 15 possible 5-taxon arrangements of the Charophyceae s. s., Coleochaetophyceae, Zygnematophyceae, Embryophyte, and Older Taxa groupings. Trees are ranked in descending order.

**Supplementary Figure 12: Principal Components Analysis of Amino Acid Frequencies in Whole Assemblies**

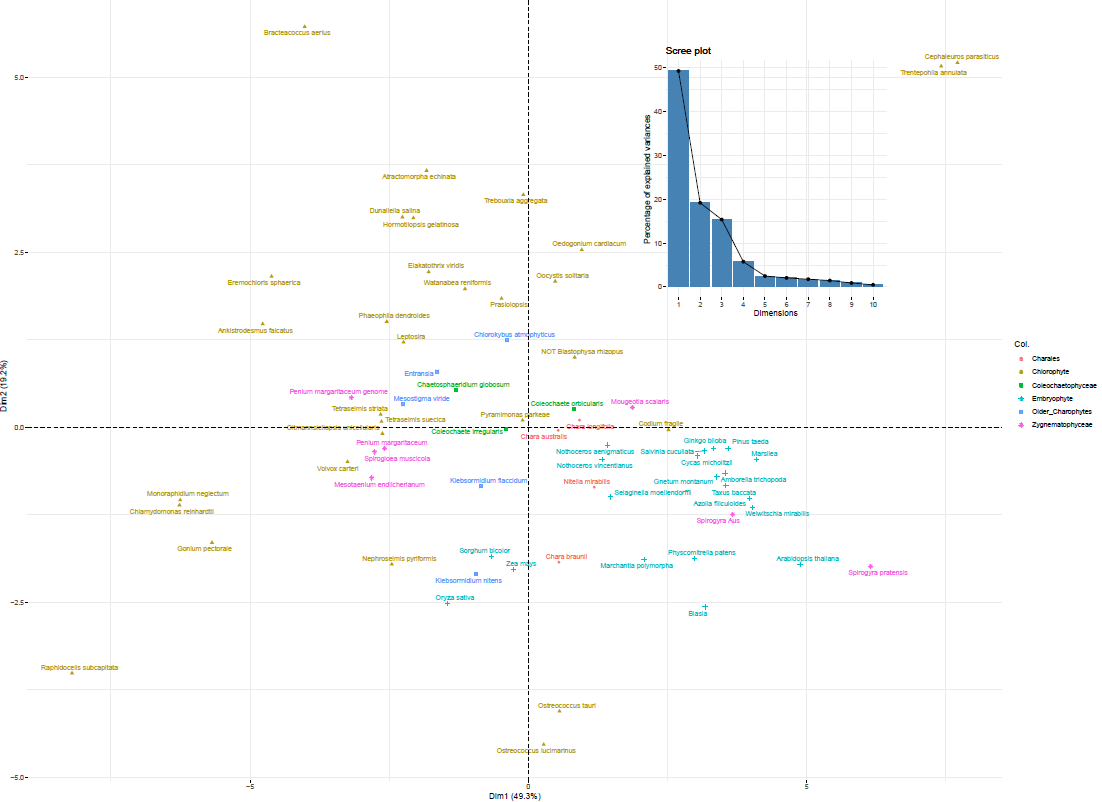

Supplementary Figure 12: Dimensions 1 and 2 of the principal components analysis of amino acid frequencies in the genomic and transcriptomic datasets. Inset scree plot shows the percentage of variation explained by each dimension.

**Supplementary Figure 13: Principal Components Analysis of Codon Frequencies in Whole Assemblies**

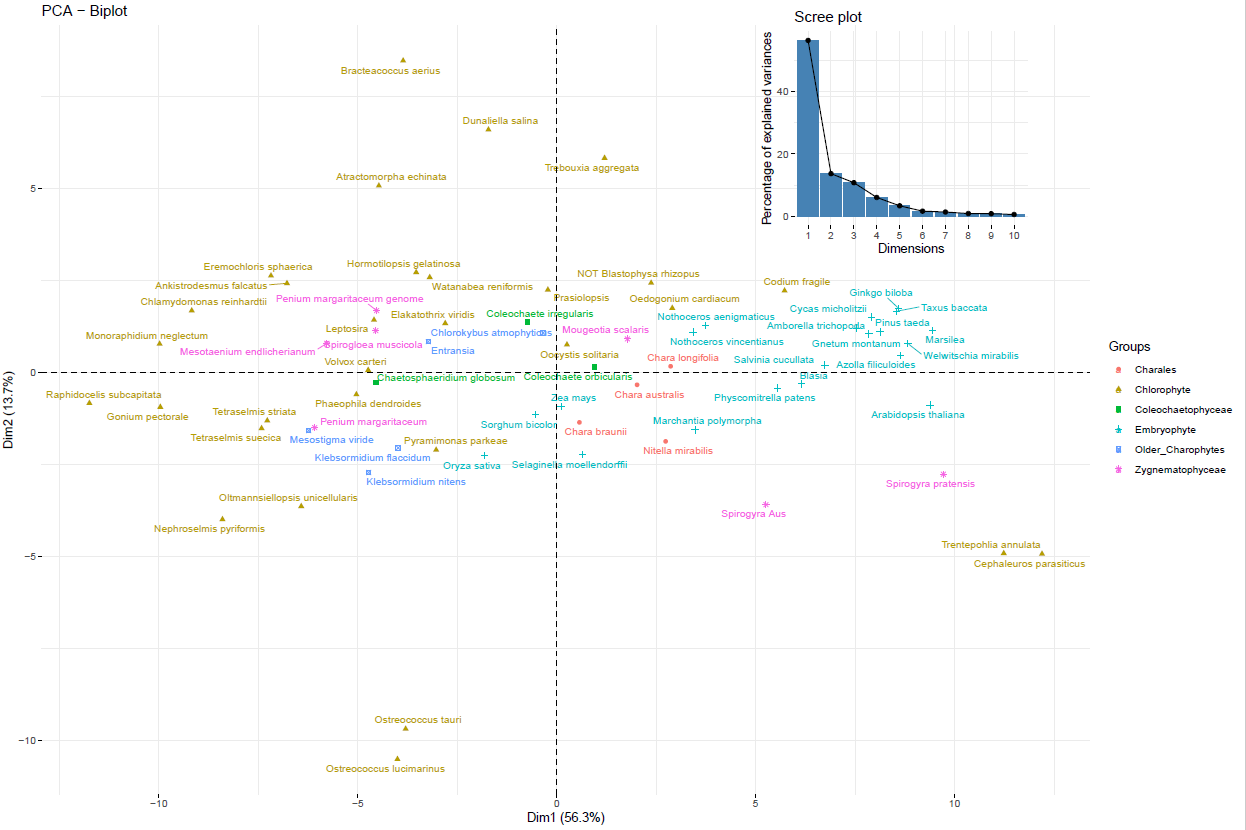

Supplementary Figure 13: Dimensions 1 and 2 of the principal components analysis of codon frequencies in the genomic and transcriptomic datasets. Inset scree plot shows the percentage of variation explained by each dimension.

**Supplementary Figure 14: ETR1 Ethylene Binding Domain Search**

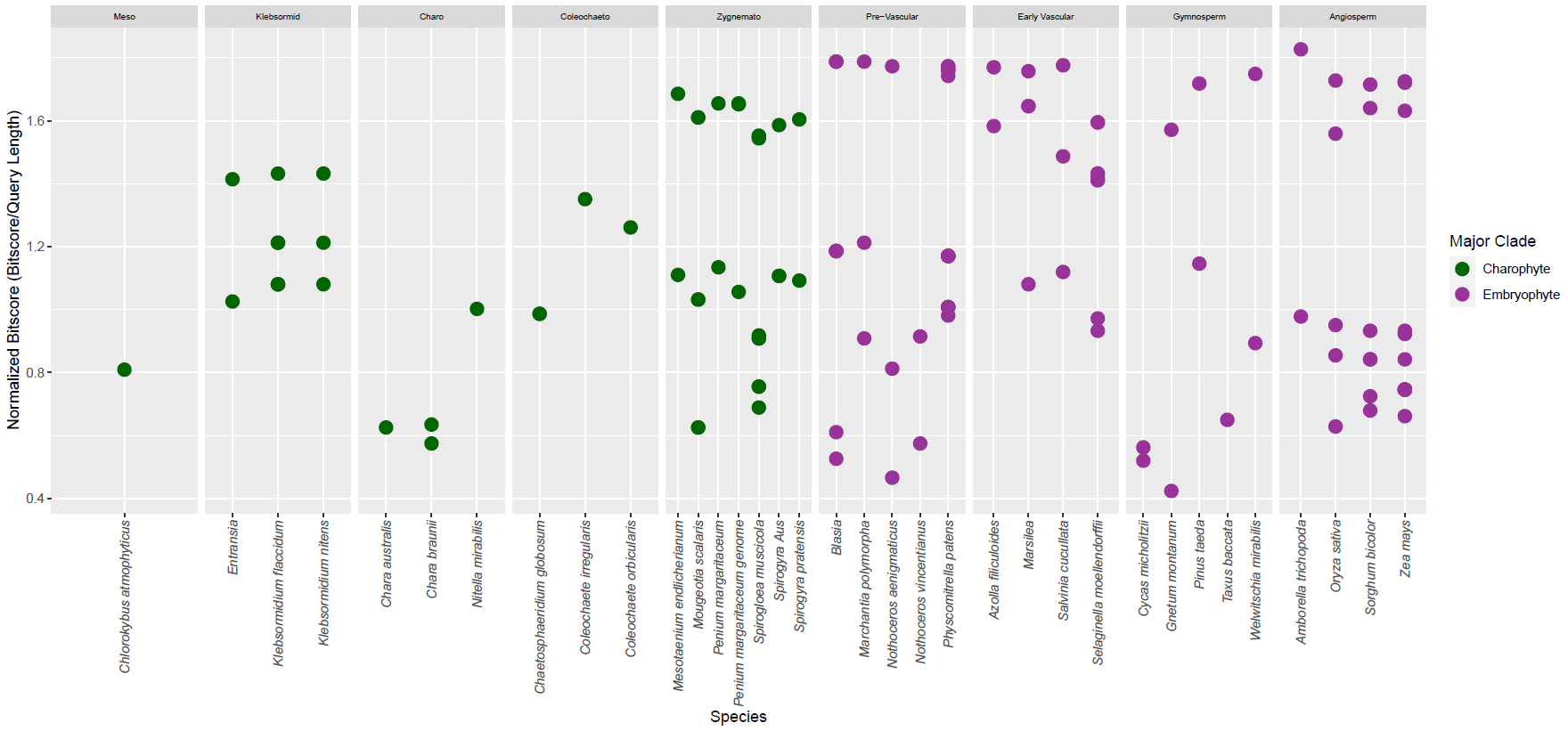

Supplementary Figure 14: BLAST search of the dataset with the ETR1 ethylene binding domain sequence of the *Arabidopsis thaliana* sequence. Clade abbreviations going from left to right: Meso=Mesostigmatophyceae, Klebsormid=Klebsormidiophyceae, Charo=Charophyceae, Coleochaeto=Coleochaetophyceae, Zygnemato=Zygnematophyceae.

**Supplementary Figure 15: EIN2 Signaling Domain Search**

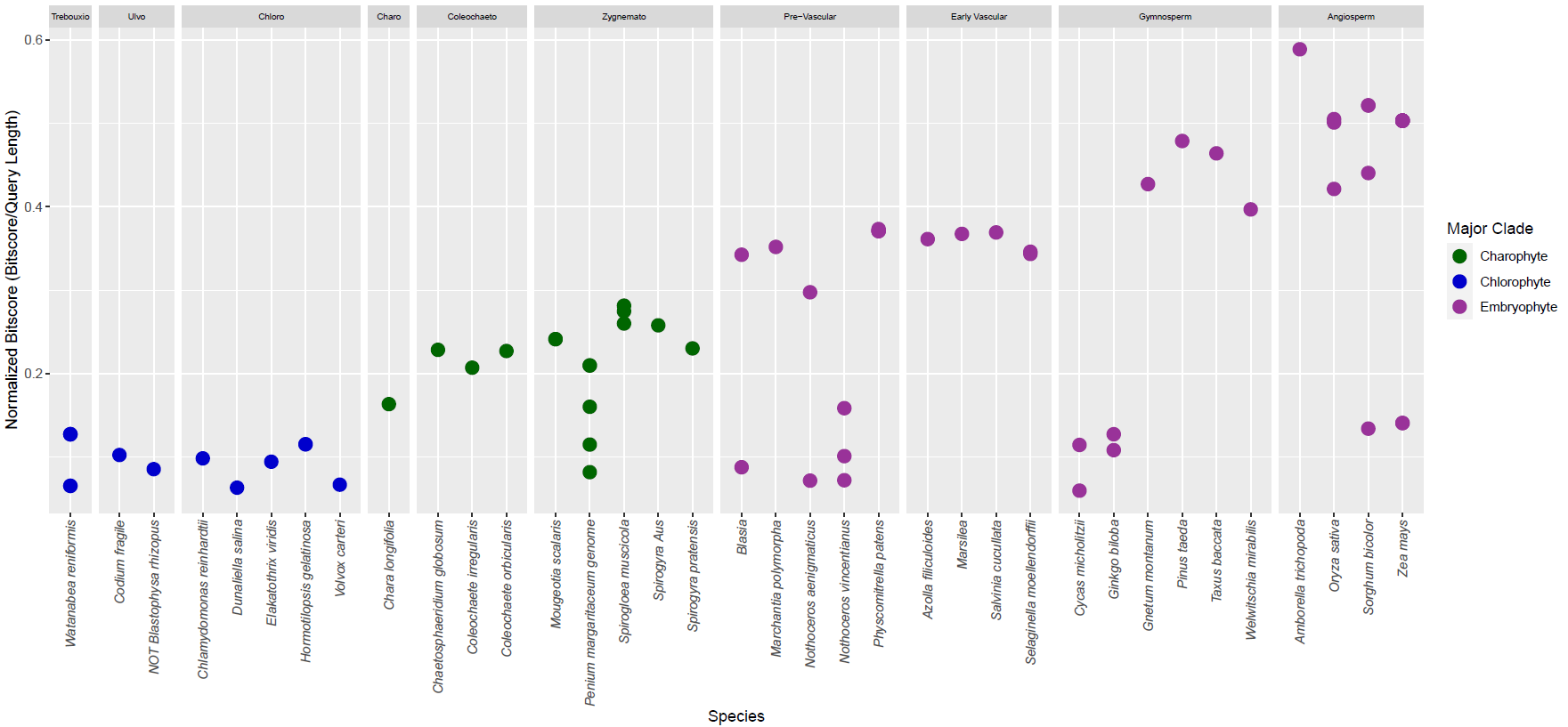

Supplementary Figure 15: BLAST search of the dataset with the EIN2 signaling domain of the *Arabidopsis thaliana* sequence. Clade abbreviations going from left to right: Trebouxio=Trebouxiophyceae, Ulvo=Ulvophyceae, Chloro=Chlorophyceae, Charo=Charophyceae, Coleochaeto=Coleochaetophyceae, Zygnemato=Zygnematophyceae.

**Supplementary Figure 16: EIN3 Gene Search**

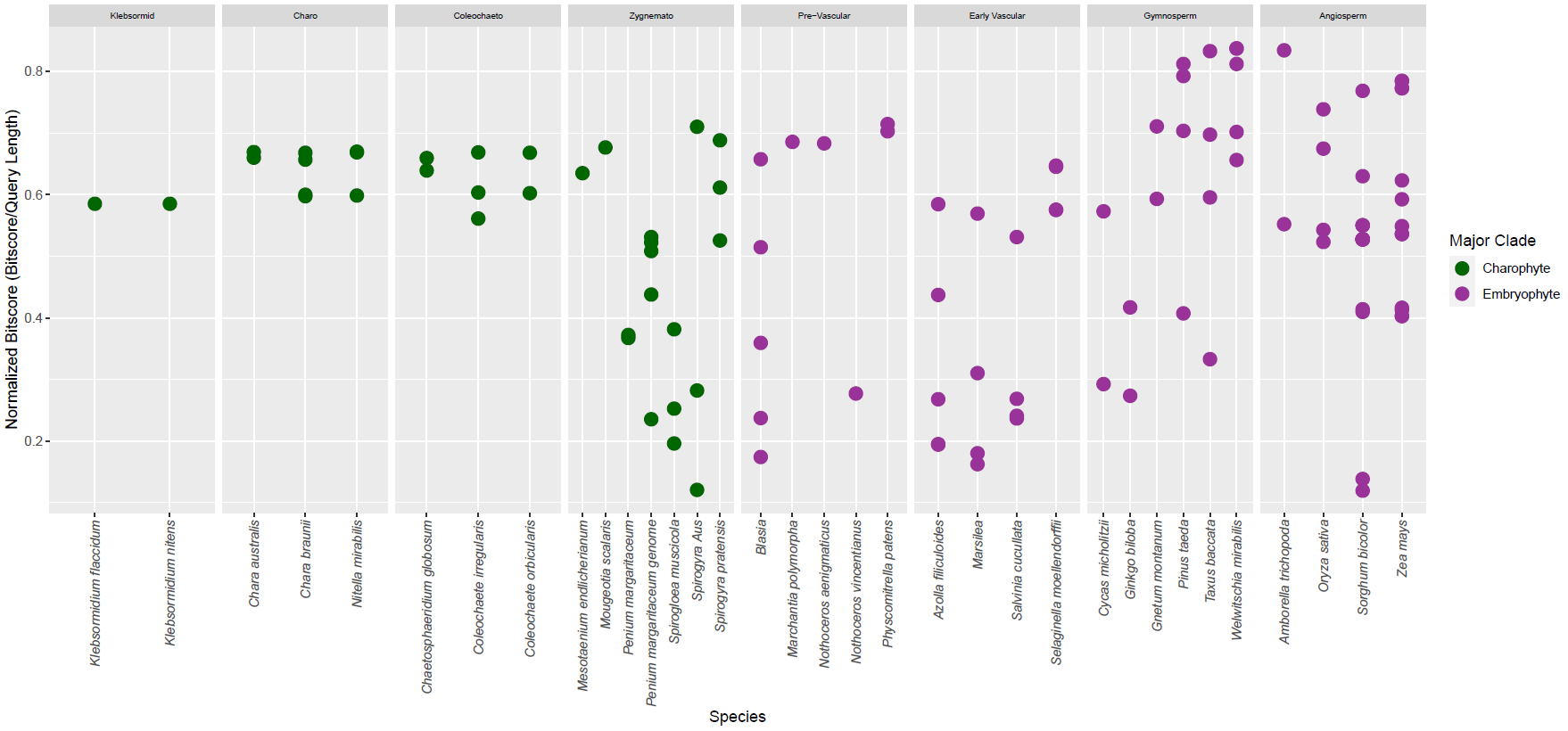

Supplementary Figure 16: BLAST search of the dataset with the EIN3 sequence with the *Arabidopsis thaliana* sequence. Clade abbreviations going from left to right: Klebsormid=Klebsormidiophyceae, Charo=Charophyceae, Coleochaeto=Coleochaetophyceae, Zygnemato=Zygnematophyceae.

**Supplementary Figure 17: CHX Gene Family Searches**

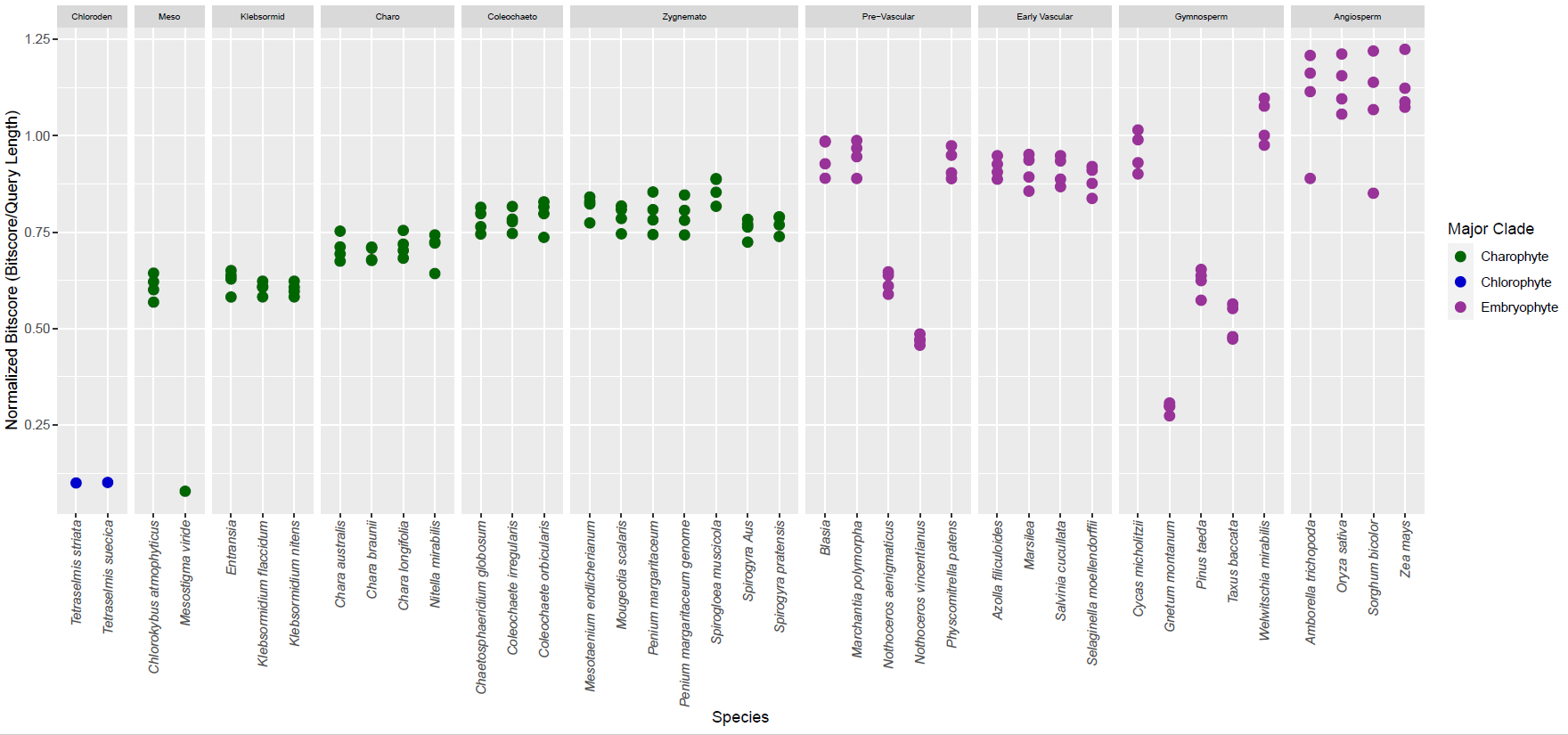

Supplementary Figure 17: BLAST search of the dataset with the CHX gene family using *Arabidopsis thaliana* sequences (n=4). To reduce clutter, only the best hit for each individual gene in the family is retained for each taxa. Clade abbreviations going from left to right: Chloroden=Chlorodendrophyceae, Meso=Mesostigmatophyceae, Klebsormid=Klebsormidiophyceae, Charo=Charophyceae, Coleochaeto=Coleochaetophyceae, Zygnemato=Zygnematophyceae.

**Supplementary Figure 18: KEA Gene Family Searches**

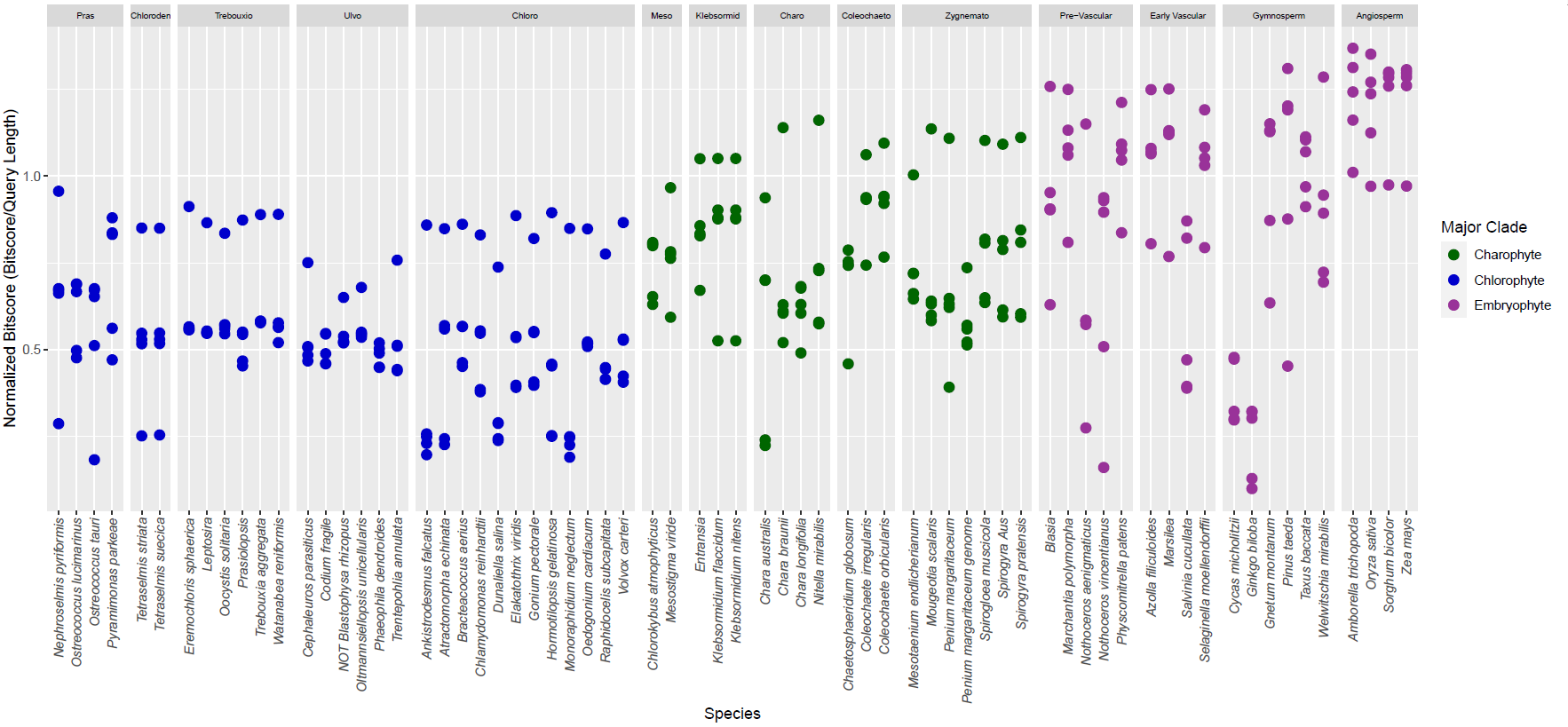

Supplementary Figure 18: BLAST search of the dataset with the KEA gene family using *Arabidopsis thaliana* sequences (n=6). To reduce clutter, only the best hit for each individual gene in the family is retained for each taxa. Clade abbreviations going from left to right: Pras=Prasinophytes, Chloroden=Chlorodendrophyceae, Trebouxio=Trebouxiophyceae, Ulvo=Ulvophyceae, Chloro=Chlorophyceae, Meso=Mesostigmatophyceae, Klebsormid=Klebsormidiophyceae, Charo=Charophyceae, Coleochaeto=Coleochaetophyceae, Zygnemato=Zygnematophyceae.

**Supplementary Figure 19: AAP Gene Family Searches**

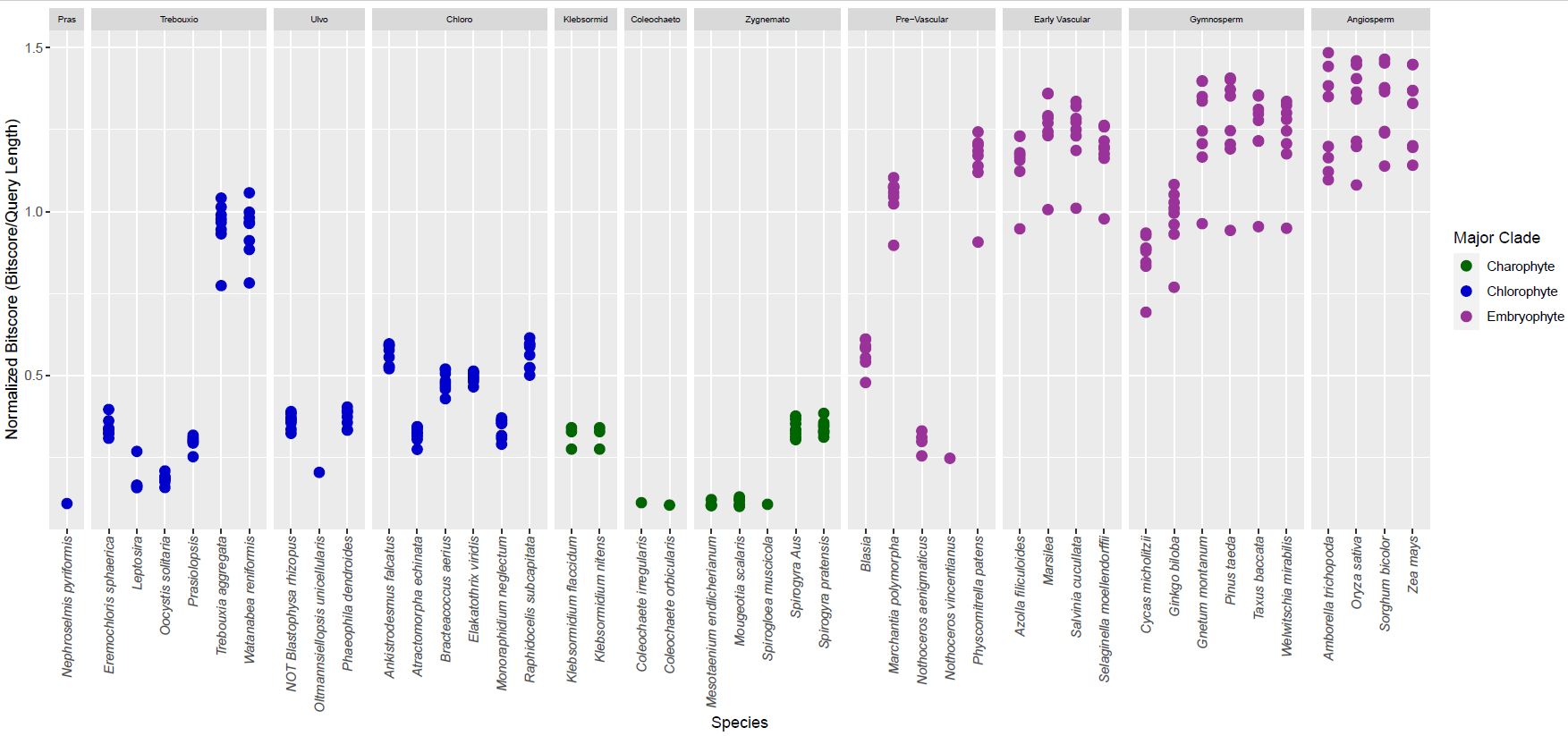

Supplementary Figure 19: BLAST search of the dataset with the AAP gene family using *Arabidopsis thaliana* sequences (n=8). To reduce clutter, only the best hit for each individual gene in the family is retained for each taxa. Clade abbreviations going from left to right: Pras=Prasinophytes, Trebouxio=Trebouxiophyceae, Ulvo=Ulvophyceae, Chloro=Chlorophyceae, Klebsormid=Klebsormidiophyceae, Charo=Charophyceae, Coleochaeto=Coleochaetophyceae, Zygnemato=Zygnematophyceae.
